# Evaluation of persistence and fate of *ex vivo* edited-HSC modified with donor template and Its role in correcting Sickle Cell Disease

**DOI:** 10.1101/2021.06.30.450644

**Authors:** Sowmya Pattabhi, Samantha N. Lotti, Mason P. Berger, Swati Singh, David J. Rawlings

## Abstract

Sickle cell disease (SCD) is caused by a single nucleotide transversion in exon 1 of the *HBB* gene that changes the hydrophobicity of adult globin (β^A^), leading to substantial morbidity and reduced lifespan. *Ex vivo* autologous gene editing utilizing co-delivery of a designer nuclease along with a DNA donor template allows for precise homology-directed repair (HDR). These gene corrected cells when engrafted into the bone marrow (BM) can prove to be therapeutic and serves as an alternative to HLA-matched BM transplantation. In the current study, we extensively explored the role of single stranded oligonucleotide (ssODN) and recombinant adeno-associated 6 (rAAV6) donor template delivery to introduce a codon-optimized change (E6optE) or a sickle mutation (E6V) change following Crispr/Cas9-mediated cleavage of *HBB* in healthy human mobilized peripheral blood stem cells (mPBSCs). We achieved efficient HDR *in vitro* in edited cells and observed robust human CD45^+^ engraftment in the BM of NBSGW mice at 16-17 weeks. Notably, recipients of ssODN-modified HSC exhibited a significantly higher proportion of HDR-modified cells within individual BM, CD34^+^ and CD235^+^ compartments of both E6optE and E6V cohorts. We further assessed key functional outcomes including RNA transcripts analysis and globin sub-type expression. Our combined findings demonstrate the capacity to achieve clinically relevant HDR *in vitro* and *in vivo* using both donor template delivery method. The use of ssODN donor template-delivery is consistently associated with higher levels of gene correction *in vivo* as demonstrated by sustained engraftment of HDR-modified HSC and erythroid progeny. Finally, the HDR-based globin protein expression was significantly higher in the E6V ssODN-modified animals compared to the rAAV6-modified animals confirming that the ssODN donor template delivery outperforms rAAV6-donor template delivery.

## Introduction

Sickle cell disease (SCD) is a hemoglobin disorder caused by a missense mutation in codon 6 (rs334) of the *HBB* gene. Patients are routinely managed through supportive care using penicillin prophylaxis, blood transfusions, hydroxyurea and iron-chelation therapy ^1-3^. HLA-matched bone marrow (BM) transplantation is a curative option for SCD patients but limited by the numbers of matched donors available in the BM registry ^4, 5^. More recently, lentiviral gene therapy that delivers an anti-sickling globin (Lentiglobin, β^T87Q^) has proved to improve quality of life in SCD patients by making them transfusion-independent ^6-9^. Other approaches that target disruption of *BCL11A* erythroid enhancer restores fetal hemoglobin expression in adult cells and is currently being tested in clinical trials with promising early results ^10^. Targeted gene editing using Crispr/Cas9 nuclease disruption followed by HDR mediated targeted integration using rAAV6 or ssODN donor template delivery remains an attractive alternative to lentiviral gene therapy, as this approach replaces the defective gene in a site-specific manner.

HDR-mediated correction has been demonstrated by various groups in both healthy cells as a proof of concept as well as in sickle patient cells. These publications demonstrate the feasibility of homology-directed repair mediated targeted correction of mutations at *HBB* ^11-17^. Our prior work directly compared rAAV6 delivery to ssODN delivery in healthy mobilized peripheral blood stem cells (mPBSCs) and demonstrated that ssODN delivery out-performed rAAV6 through sustained engraftment of ssODN-modified cells in the BM at 12-14 weeks in NBSGW mouse model ^17^. The testing was done using high-density culturing and Neon electroporation system in healthy cells to test the introduction of the sickle mutation at *HBB*.

In our current work, we have tested the role of donor template delivery in achieving therapeutic levels of engraftment using widely accepted enhancements that improves and preserves long term hematopoietic stem cells (LT-HSCs) using low-density culturing with UM171 ^18^ + SR-1 ^19^ (StemRegenin 1) and Lonza nucleofection system. As a proof of concept, we tested both the introduction of the sickle mutation (E6V GTC, glutamate to valine amino acid change) along with the introduction of a codon-optimized change (E6optE GAA, glutamate to glutamate) into healthy mPBSCs. The *in vitro* HDR achieved was compared to the *in vivo* maintenance of gene-edited cells in the BM, CD34^+^- and CD235^+^- enriched compartments at 16-17 weeks in NBSGW mouse model ^20^. We further evaluated the RNA transcript levels and β-like globin protein expression produced by the HDR integration by measuring the alteration of gene expression in the *HBB* and *HBG* gene in individual BMs as well as in the CD34^+^- and CD235^+^- enriched BM pools at the time of sacrifice. Consistently the ssODN donor template delivery outperformed rAAV6 donor template delivery through the higher frequency of gene-modified cells in the BM which results in higher HDR-based RNA transcript levels and concomitantly higher HDR-based globin expression in both the E6optE and the E6V cohorts of ssODN-modified animals at 16-17 weeks. This increase in gene-modified cells is also observed in the HSC and erythroid enriched compartments of the BM as well as in the spleen of the ssODN-modified animals.

## Results

### Precise gene editing with both rAAV6 and ssODN donor template delivery in adult healthy human mobilized peripheral blood CD34^+^ donors

The sgRNA was designed to disrupt at the sickle mutation site within the exon 1 of *HBB* gene. Both the ssODN and the rAAV6 donor template were designed as previously reported ^17^ to deliver either a E6optE GAA change that introduces a codon-optimized GAG to GAA change (no change in amino acid) at codon 6 of exon 1 or a E6V GTC change that introduces a GAG to GTC (glutamate to valine change), which introduces a sickle mutation into healthy mPBSCs (Figure 1A) ^13^. The nuclease cleavage resulted in double-stranded breaks adjacent to codon 6 of exon 1. The overall NHEJ with the RNP alone was an average of 83.5 ± 7% (n = 4 transplants, 2 CD34^+^ donors). The average residual NHEJ remaining after HDR in the E6optE ssODN- and rAAV6-modified input cells were 38.8 ± 11.4% and 52 ± 12.4% respectively and in the E6V ssODN- and rAAV6-modified input cells were 40.8 ± 12.1% and 31 ± 0% respectively (Figure 1B). The NHEJ was further measured across all groups on day 5, 9 and 14. A slight increase in average % NHEJ was observed from day 5 to day 14 (Figure S1A). The overall viability was slightly lower for ssODN-edited (E6optE 77.9 ± 12.1% and E6V 86.7± 3.4%) on day 1 post-editing compared to the rAAV6-edited groups (E6optE 90.7 ± 3% and E6V 93.2 ± 1.7%, Figure S1B).

**Figure 1.**
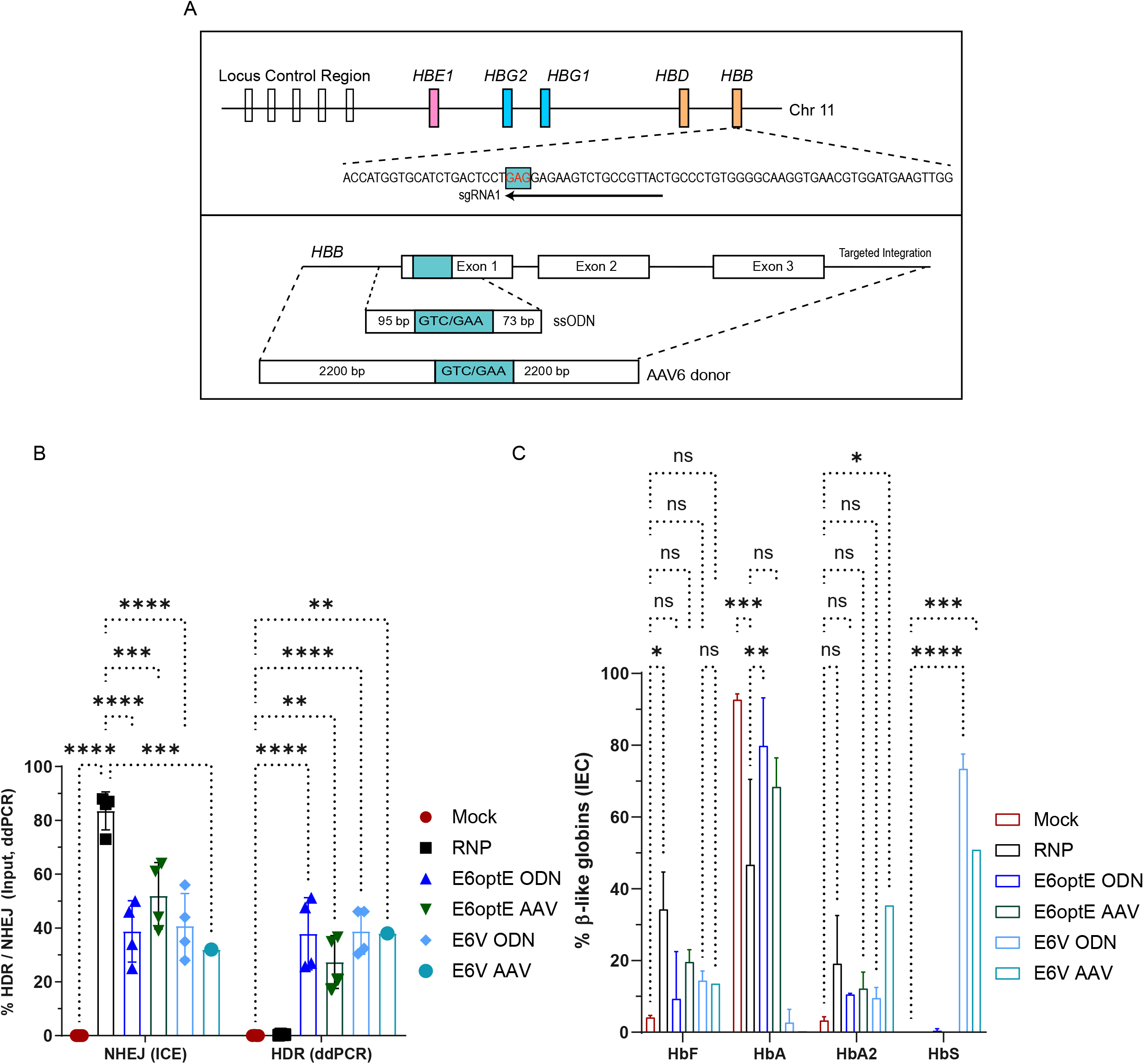
HDR donor template design and gene editing at codon 6 of *HBB*. (A) The structural organization of β-globin locus and *HBB* on Chromosome 11. The sgRNA was designed to cut at the site of sickle mutation shown in red. The ssODN donor template was 168 bp long and the rAAV6 donor template was designed with a 2.2 Kb homology repair (HR)-arm on both the 5’ and 3’ ends. Both donor templates introduce either a E6optE GAA (no change in amino acid) or a E6V GTC (Glutamate to Valine change) at codon 6 of exon 1 of *HBB*. (B) The NHEJ % measured by ICE algorithm and HDR % measured by ddPCR using a dual probe assay (FAM/HEX) in *in vitro* cultured input cells on day 14 post-editing (n = 4 transplants, 2 different CD34+ donors). (C) Ion-exchange chromatography (IEC) of *in vitro* differentiated erythroid cells to quantify globin tetramers: adult (HbA), fetal (HbF), minor adult (HbA2) and sickle (HbS). Statistical analyses were run using 2-way ANOVA using Tukey’s multiple comparison test (P-value of <0.0001, *** < 0.001, ** <0.01).

An average HDR of 37.9 ± 13.4% and 27.4 ± 9.8% was achieved with the ssODN- and rAAV6-edited respectively introducing the E6optE GAA change and an average HDR of 38.8 ± 8.5%, 38 ± 0% was achieved with ssODN- and rAAV6-edited input respectively introducing the E6V GTC change (n = 4 transplants with 2 CD34^+^ donors, except RNP + GTC AAV: n = 1 transplant, Figure 1B). The % HDR measured by ddPCR on day 10 and 14 were comparable across all groups (Figure S1C).

The β-like globin sub-types were measured in differentiated erythroid cells on day 14 post-editing by ion-exchange chromatography (IEC). There was a decrease in HbA levels (Mock: 92.7 ± 1.7%, RNP: 46.7 ± 23.8%) and an increase in HbF levels (Mock: 4.1 ± 0.6%, RNP: 34.2 ± 10.4%) in the RNP-edited cells compared to the mock-edited cells. On comparing HbA levels of RNP alone (46.7 ± 23.8%) with the E6optE GAA change, significantly higher levels of restoration of HbA was observed with the ssODN-modified cells (79.8 ± 13.4%) compared to the rAAV6-modified cells (68.4 ± 8.1%). Similarly, the E6V GTC change had higher level of sickle globin protein expression with the ssODN-edited cells (73.4 ± 4.1%) compared to the rAAV6-edited cells (50.9 ± 0%, Figure 1C).

#### *In vivo* engraftment of gene modified cells in the BM

Human adult mPBSCs mock-edited, RNP-edited, ssODN-modified and rAAV6-modified introducing E6optE GAA or E6V GTC change were transplanted into NBSGW mice (n = 4 transplants, 2 CD34^+^ donors, Figure 2A). Robust human cell engraftment was observed in all cohorts. Mock-edited and RNP-edited cohorts had an average human CD45^+^ (hCD45^+^) engraftment of 76.1 ± 14.4% and 81.8 ± 4.9% respectively. Though not significant, a 1.15 - 1.33-fold decline in average hCD45^+^ was observed in the edited cohorts modified with donor template delivery (Figure 2B).

**Figure 2.**
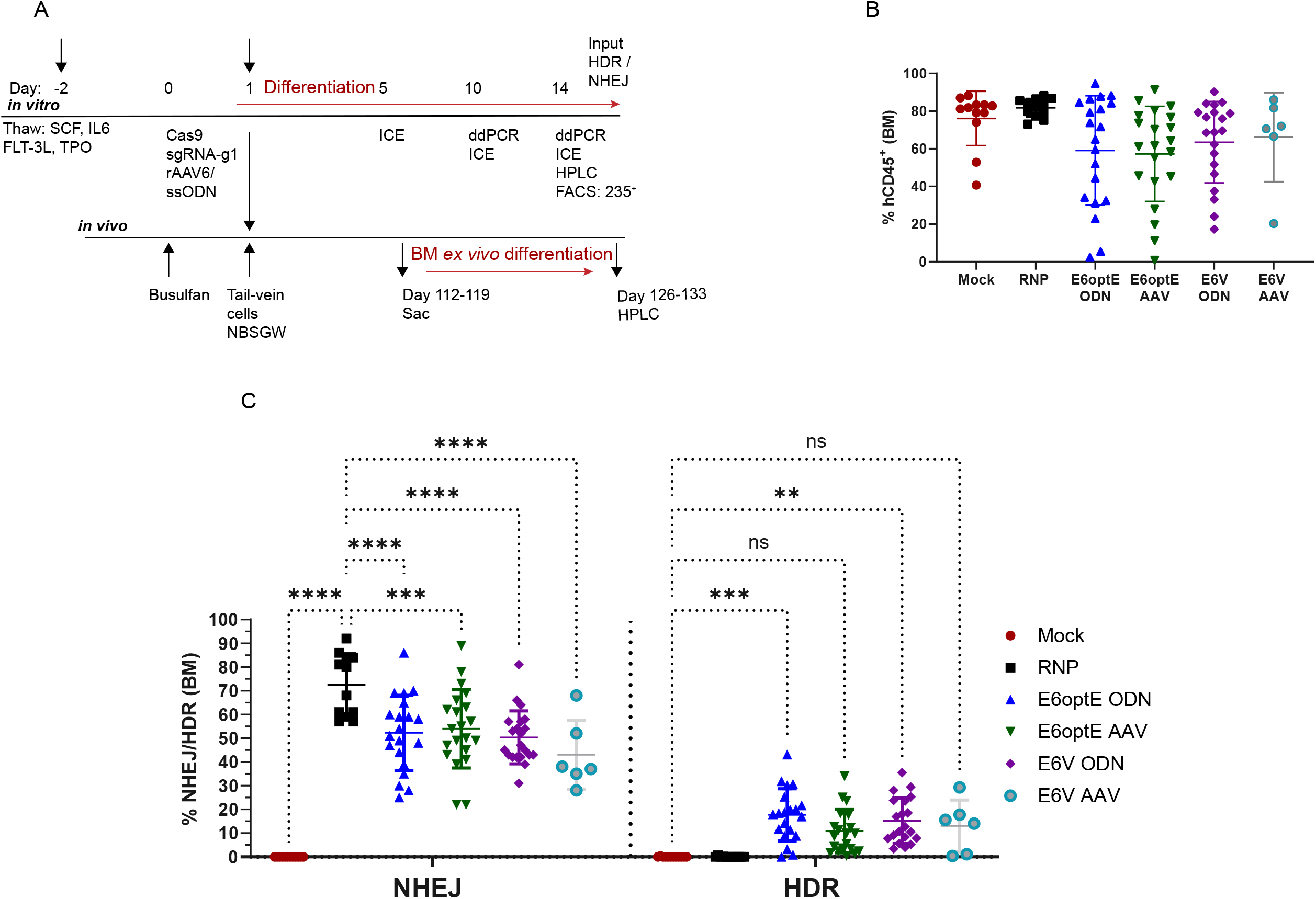
*In vivo* engraftment of gene-edited cells in the BM of NBSGW mice at 16-17 weeks. (A) Schematic of the experimental details for both *in vitro* and *in vivo* studies. (B) Human chimerism (hCD45^+^) measured in the BM at 16-17 weeks across mock-edited (n = 12 animals), RNP-edited (n = 12 animals), E6optE ssODN (n = 20 animals) / rAAV6-modified (n = 21 animals), E6V ssODN (n = 20 animals) / rAAV6-modified (n = 6 animals) cohorts. Each dot represents a unique animal in the graph (C) NHEJ % measured using ICE algorithm and HDR % measured by ddPCR across various cohorts in the BM at individual animals at 16-17 weeks.

The average NHEJ measured in the BM of the RNP-edited cohort was 73.5 ± 13.2% (n= 12 animals). The residual NHEJ in donor template modified groups were 52.3 ± 15.9% and 54 ± 16.6% in the ssODN and rAAV6-modified cohorts respectively introducing the E6optE GAA change and 50.4 ± 11.2% and 43 ± 14.5% in the ssODN and rAAV6-modified cohorts respectively introducing the E6V GTC change (Figure 2C). The NHEJ alleles in the ssODN and rAAV6-edited cohorts showed a 1.03 – 1.35-fold average increase in the BM at 16-17 weeks compared to the pre-transplant levels.

The average HDR-modified cells measured in the BM at 16-17 weeks was 17.7 ± 11% and 10.8 ± 9.1% for ssODN (n = 4 transplants, 20 animals) and rAAV6 (n = 21 animals) delivery respectively introducing the E6optE GAA change and 15.2 ± 9.6% and 13 ± 10.9% for ssODN (n = 21 animals) and rAAV6 (n= 6 animals) delivery respectively introducing the E6V GTC change (Figure 2C, S2). The fold decrease in average HDR-modified cells compared to the input HDR was more pronounced with the rAAV6-modified cells (E6optE: 2.53-fold, E6V: 2.91-fold) in the BM than the ssODN-modified cells (E6optE: 2.14-fold, E6V: 2.54-fold, Figure 2C).

All cohorts showed multi-lineage engraftment with comparable levels of CD34^+^, CD34^+^ 38^lo^, CD235^+^, CD33^+^, CD19^+^ in the BM (Figure S3A-F, S4). Though not significant, the average myeloid populations were lower in the rAAV6 modified cohorts (E6optE: 36.5 ± 10.4% and E6V: 29.2 ± 7.3%) compared to the ssODN modified cohorts (E6optE: 41.6 ± 9.7% and E6V: 40.7 ± 9.4%, Figure S3E). The average lymphoid populations were lower in the ssODN modified cohorts (E6optE: 48.6 ± 10.2 % and E6V: 49.9 ± 10.6%) than the rAAV6 modified cohorts (E6optE: 53.5 ± 11.7 % and E6V: 62.3 ± 7.4 %, Figure S3F).

#### *In vivo* engraftment of gene corrected cells in the CD34^+^, CD235^+^ enriched BM pools and the spleen

The cells isolated from the BM of animals were pooled within each cohort and the HSC (CD34^+^) and erythroid (CD235^+^) compartments were enriched using magnetic beads to measure the level of gene-modified cells in each of these compartments. A ∼ 5.7-fold enrichment was achieved with the CD34^+^ compartment from BM pools (Figure S5A, n = 3 transplants). The average % HDR-modified cells in the CD34^+^ enriched compartment of ssODN-modified cohorts were higher than the rAAV6-modified cohorts (E6optE CD34^+^: ssODN: 25.1 ± 13% and rAAV6: 9.6 ± 4.7%; E6V CD34^+^: ssODN:15.8 ± 7.8% and rAAV6: 8.2 ± 0%, Figure 3A).

**Figure 3.**
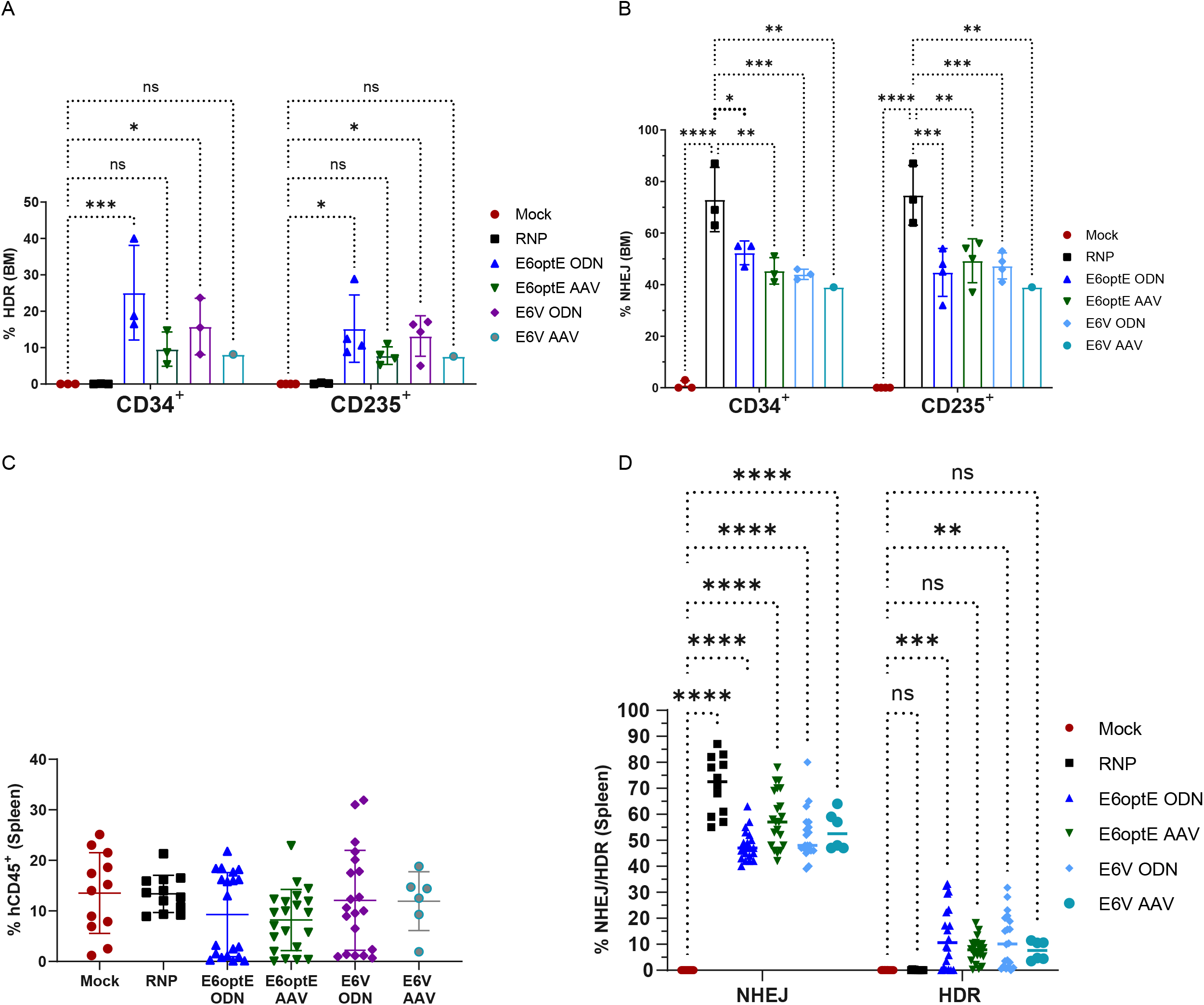
*In vivo* HDR and NHEJ in the hCD34^+^, hCD235^+^ enriched compartments and the spleen. (A) HDR% measured in hCD34^+^ and hCD235^+^ enriched compartments from BM pools at 16-17 weeks by ddPCR across various cohorts. (B) NHEJ% measured in hCD34^+^ and hCD235^+^ enriched compartments from BM pools at 16-17 weeks using ICE algorithm across various cohorts. (C) Human chimerism (hCD45^+^) measured in the spleen across mock-edited, RNP-edited, E6optE ssODN/rAAV6-modified, E6V ssODN/rAAV6-modified cohorts. (D) NHEJ% measured by ICE algorithm and HDR% measured by ddPCR in the spleen of individual animals across various cohorts. Statistical analyses were run using 2-way ANOVA using Tukey’s multiple comparison test (P-value of <0.0001, *** < 0.001, ** <0.01).

Further, the CD235^+^ compartment was enriched from the flow through fraction containing the CD34^-^ fraction. A 11-fold increase in CD235^+^ population was achieved in the enriched fractions (Data not shown). The average % HDR-modified cells in the CD235^+^ enriched compartment of ssODN-modified cohorts were higher than the rAAV6-modified cohorts (E6optE CD235^+^: ssODN: 15.2 ± 9.3% and rAAV6: 7.8 ± 2.4%; E6V CD235^+^: ssODN: 13.2 ± 5.6% and rAAV6: 7.6 ±.0%, Figure 3A).

The % NHEJ in the CD34^+^ purified compartment was comparable to the % NHEJ observed in the CD235^+^ enriched compartment (E6optE CD34^+^ ssODN: 52.3 ± 4.6% and rAAV6: 45.3 ± 5.1%; E6V CD34^+^ ssODN: 44 ± 2% and rAAV6: 39 ± 0%; E6optE CD235+ ssODN: 44.8 ± 9.3% and rAAV6: 49.3 ± 8.5%; E6V CD235^+^ ssODN: 47.3 ± 5.1% and rAAV6: 39 ± 0%, Figure 3B).

The ability of the edited HSCs to repopulate a secondary lymphoid organ was evaluated by measuring the human chimerism in spleen. The hCD45^+^ engraftment in the spleen was comparable across groups. Mock-edited, RNP-edited had an average engraftment of 13.5 ± 8% and 13.4 ± 3.7% respectively. The average hCD45^+^ engraftment of the edited cohorts was slightly lower (average 10.4 ± 1.9%) than the mock-edited cohort. ssODN-modified and rAAV6-modified cohorts introducing E6optE change had 9.3 ± 8.3% and 8.2 ± 6.1% hCD45^+^ respectively and the ssODN-modified and rAAV6-modified cohorts introducing E6V change had hCD45^+^ engraftment of 12.1 ± 9.9% and 11.9 ± 5.8% respectively (Figure 3C).

The average NHEJ measured in RNP-modified animals were 71.2 ± 11.1% (n = 12 animals). The average residual NHEJ measured in the ssODN- and rAAV6-modified animals introducing E6optE change was 47.9 ± 5.8% and 58.2 ± 10.7% respectively and E6V change was 51.9 ± 9.4% and 53.7 ± 7.3% respectively (Figure 3D). The average HDR-modified cells measured in the spleen by ddPCR was 13.1 ± 12% and 7.6 ± 4.7% in the ssODN-modified and rAAV6-modified cohorts respectively introducing E6optE GAA change and 11.3 ± 10.2% and 7.5 ± 3.7% in the ssODN-modified and rAAV6-modified respectively introducing E6V GTC change (Figure 3D).

The CD33^+^ and CD19^+^ populations were comparable across groups in the spleen. The myeloid populations were lower in the rAAV6 cohorts (E6optE: 11% and E6V: 8.8%) compared to the ssODN cohorts (E6optE 15.2% and E6V 15.6%, Figure S5B). Though not significant, the average lymphoid populations were lower in the ssODN cohorts (E6optE: 75.4% and E6V 78.4%) compared to the rAAV6 cohorts (E6optE 78.4% and E6V 86.4%, Figure S5C).

#### Modulation of Globin RNA and hemoglobin in the BM

The transcripts generated by the HDR event along with the WT *HBB* and *HBG* RNA were quantified in the BM of animals at the time of sac to measure the perturbance of β-like globin expression following gene editing using a ddPCR assay (Figure S6). *GAPDH* RNA was used as a reference to compare across samples. The HDR RNA levels measured in the BM was compared to the respective globin protein in ex vivo differentiated BM cultures to evaluate the functional modulation of hemoglobin achieved through editing and gene modification. A subset of all the engrafted animals within each cohort that had a positive readout for BM HDR (%) were compared to understand the functional modulation of globin RNA and/ globin protein expression in the BM.

#### Functional outcome of Cas9 RNP editing

The NHEJ alleles were well preserved in the BM of RNP-edited animals (63 ± 23%) at 16 weeks. The nuclease cleavage increased average HBG RNA levels in the RNP-edited (17.7 ± 7.9%, n = 12 animals) compared to the mock-edited animals (2.5 ± 2.1%, n = 9 animals). This corresponded to an average HbF protein of 20.6 ± 7.9% in the RNP-edited animals compared to 8.5 ± 8.4% HbF in the mock-edited animals (Figure S7 A-B).

The WT alleles that were measured in the BM were between 17.4 - 31.1 % across the individual animals in various groups. This resulted in a reduction in average WT HBB RNA levels in the RNP-edited cohort (82.2 ± 7.9%) compared to the Mock-edited cohort (97.4 ± 2%) and resulted in a reduction of average levels of HbA protein in the RNP-edited (46.1 ± 14%, n =12 animals) compared to the Mock-edited animals (84.8 ± 11.2%, n = 9 animals, Figure S7C-D).

#### Functional outcome of E6optE GAA change

The % average HDR measured by ddPCR in the BM although not significant, was higher for ssODN-modified animals (20.5 ± 11.2%, n = 15 animals) compared to the rAAV6-modified animals (11.3 ± 9.9%, n = 15 animals) and this correlated to relatively higher average HDR RNA transcript levels for ssODN-modified animals (34.1 ± 19.8 %, n = 15 animals) compared to the rAAV6-modified animals (24.6 ± 24.6%, n = 15 animals). The RNA transcripts were measured using a sequence specific probe assay and uniquely identifies the HDR (GAA) vs the WT (GAG) transcripts (Figure S6), whereas the HbA protein levels are non-descript as both WT (GAG) and the HDR (GAA) change encodes a glutamate in the 6^th^ position of exon 1 and results in HbA expression. The HbA protein levels were further altered with a trend towards higher average HbA protein levels (58.3 ± 18.3%, n = 13 animals) for ssODN-modified animals compared to the rAAV6 modified animals (53.4 ± 29.2%, n = 12 animals, Figure 4A, 4C).

**Figure 4.**
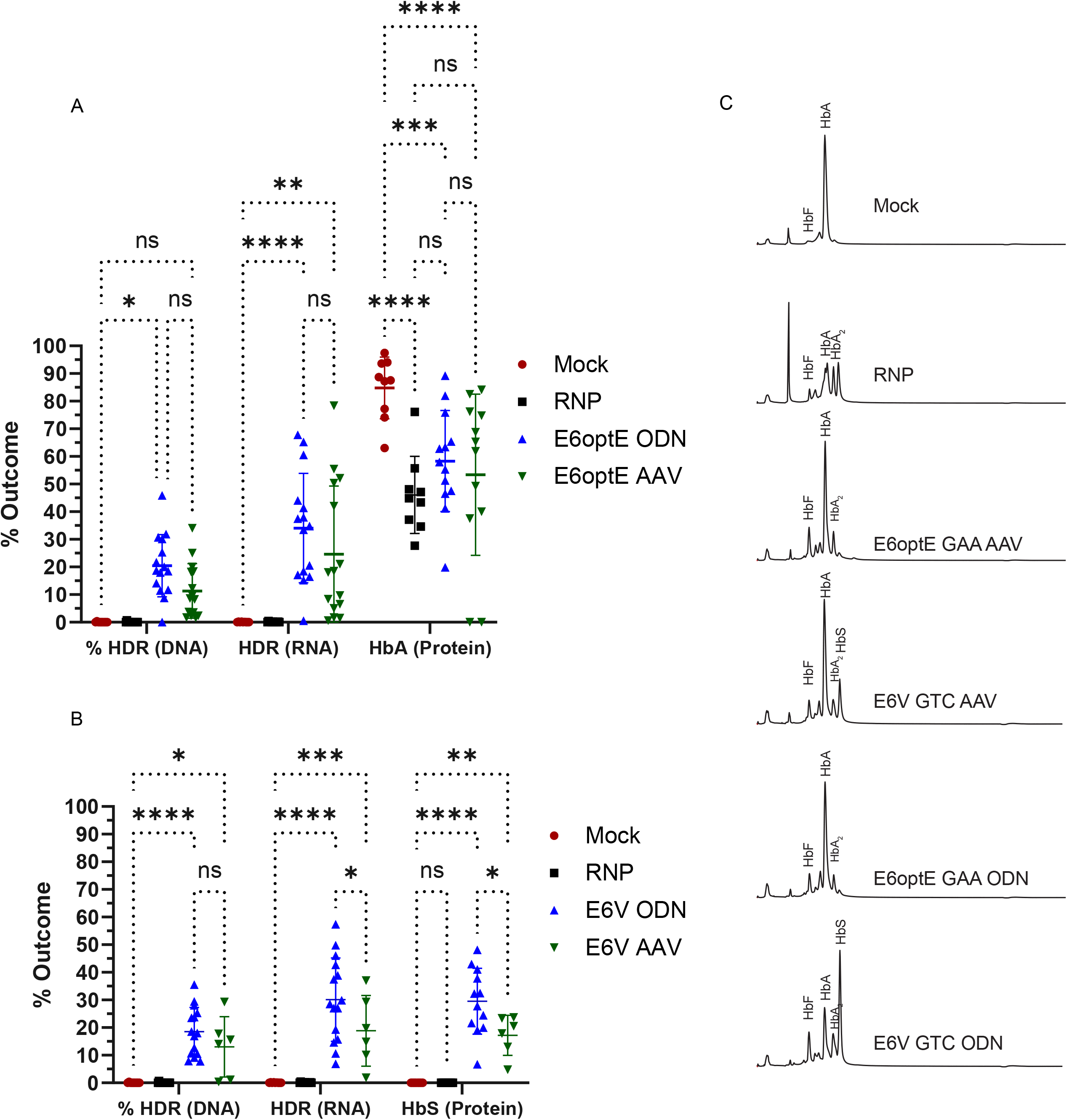
*In vivo* HDR, RNA in the BM of NBSGW mice at 16-17 weeks and globin protein analysis in *ex vivo* erythroid differentiated BM cultures. (A) BM HDR and HDR RNA transcript levels measured in individual animals and HbA protein levels measured in the individual *ex vivo* erythroid differentiated BM cultures of (A) E6optE ssODN/rAAV6 cohorts (B) E6V ssODN/rAAV6 cohorts. (C) Representative IEC trace from ex *vivo* differentiated erythroid cells from individual BM of mock-edited, RNP-edited, E6optE ssODN/rAAV6-modified, E6V ssODN/rAAV6-modified animal to quantify globin tetramers namely; adult (HbA), fetal (HbF), minor adult (HbA2) and sickle (HbS). Statistical analyses were run using 2-way ANOVA using Tukey’s multiple comparison test (P-value of <0.0001, *** < 0.001, ** <0.01).

The residual NHEJ in the E6optE ssODN-modified cohort (52.4 ± 13.1%, n = 15 animals) was comparable to the rAAV6-modified animals (56.1 ± 15.7%, n = 15 animals). This resulted in comparable levels of average HBG RNA in ssODN-modified animals (11.5 ± 10.3%) and rAAV6-modified animals (11.7 ± 8.6%). The average HbF protein expression was slightly lower in the ssODN-modified animals (21.7 ± 12.4%) compared to rAAV-modified animals (31.5 ± 34.3%, Figure S7A).

The average %WT was comparable across E6optE ssODN (18.4 ± 7.9%) and rAAV6 (17.4 ± 10.7%) modified animals and resulted in WT HBB average RNA levels that were comparable (ssODN:54.5 ± 16.7%, rAAV6: 63.7 ± 23.3%). The average HbA protein expression that does not discriminate between the HDR and WT alleles had a trend towards higher HbA protein in the ssODN modified animals (58 ± 18.3%) compared to the rAAV6-modified animals (53.4 ± 29.2%, Figure S7C).

#### Functional outcome of E6V GTC change

The average BM HDR, although not significant, were higher in the ssODN modified cohort (18.5 ± 8.8%, n= 15 animals) compared to the rAAV6-modified cohort (13.0 ± 10.9%, n = 6 animals). This resulted in a significantly higher average HDR RNA level for the ssODN-modified animals (30.1 ± 15.1%, n = 15 animals) compared to the rAAV6-modified animals (18.8 ± 12.8%, n = 6 animals). The HbS levels measured in ex vivo differentiated erythroid cultures from the BM had significantly higher average HbS level in the ssODN-modified animals (29.5 ± 11.9%, n = 12 animals) compared to the rAAV6-modified animals (17.2 ± 7.2%, n = 6 animals, Figure 4B-C).

The residual NHEJ in the BM was similar for ssODN-modified animals (49.8 ± 11.2%, n = 15 animals) and the rAAV6-modified animals (43 ± 14.5%, n = 6 animals). This resulted in comparable levels of HBG RNA in the ssODN cohort (11.7 ± 10.1%, n = 15 animals) and the rAAV6-modifed cohort (6.04 ± 4.3%, n = 6 animals). Though not significant, the average HbF expression almost doubled in the ssODN animals (24.2 ± 11.6%, n = 15 animals) compared to the rAAV6-modified animals (12.7 ± 8.3%, n = 6 animals, Figure S7B).

The average %WT alleles measured were comparable in the ssODN- and rAAV6-modified cohorts (ssODN:19.5% ± 8.3%, rAAV6: 31.1% ± 11.8%). This resulted in a concomitant decrease in WT RNA and HbA protein levels compared to the Mock-edited animals (Figure S7D).

### Functional outcome CD34^+^ and CD235^+^ enriched compartments

The HDR, NHEJ and WT alleles were measured in the CD34^+^ and CD235^+^ enriched compartments of BM pools. The HDR in the BM pools had a trend towards higher HDR in the ssODN-modified animals compared to the rAAV6 modified animals of both E6optE and E6V cohorts (Figure S5A-D). This resulted in a 1.87 - 2.4-fold increase in average HDR RNA in the ssODN-modified cohort compared to the rAAV6 cohort of the CD34^+^ and CD235^+^ enriched compartments of both E6optE and E6V groups. The ssODN-modified animals had higher average HbA expression in the BM pools of E6optE cohorts and significantly higher HbS expression in the BM pools of E6V cohorts (Figure S8A-D).

The NHEJ alleles were persistent in the CD34^+^ and CD235^+^ enriched compartments of the BM pools at 16-17 weeks and there was and trend towards an increase in NHEJ alleles in the BM pools compared to the input (S9A-B). The persistent NHEJ in the BM in the edited cohorts resulted in an average increase in % *HBG* RNA and a decrease in % *HBB* RNA compared to the Mock-edited animals (Figure S9A-D). The HbF protein was elevated in all the edited cohorts compared to the Mock-edited animals (Figure S9A-B).

The WT alleles measured in the BM enriched pools of the edited cohorts were between 24.2 - 33.5% across donor template modified groups. This resulted in a 1.6-fold reduction in average WT-specific HBB RNA levels in the edited cohorts compared to the Mock-edited cohort (97.4 ± 2%) and resulted in a 1.9-fold in reduction of HbA protein compared to the Mock-edited animals (Figure S9C-D).

## Discussion

Autologous gene editing can serve to be an alternative to HLA-matched allogeneic BM transplant and lentiviral gene therapy in SCD patients ^7-9^. A targeted gene replacement approach proves to be a safer option than semi-random integration though lentiviral gene therapy ^21^, as it allows for site-specific integration of the corrected gene through template-driven repair following site-specific nuclease disruption. This allows for the correction at the site of mutation and preserves the normal regulation and intrinsic controls that allow for normal levels of protein expression. The sickle mutation in exon 1 (rs334) of *HBB* can be efficiently corrected through targeted nuclease disruption and homology-directed repair in autologous cells ^11-16^

Numerous studies have suggested that a 10-20% mixed chimerism is sufficient to alleviate SCD complications, increase hemoglobin levels and reduce HbS%. Even with low donor chimerism, the selective advantage of healthy RBCs is sufficient to replace sickle RBCs and therefore proves to be therapeutic ^5, 22-24^. Clinical studies have also shown that ∼ 10 - 20% restoration of fetal globin can alleviate major organ damage as well as pulmonary complications in SCD ^25, 26^. We were interested in understanding the role of donor templates in driving HDR in mPBSCs, understanding the long-term fate of *ex vivo* edited mPBSCs and their persistence in the BM of NBSGW mice at 16-17 weeks. We used viral-mediated rAAV6 delivery and compared to non-viral ssODN delivery to evaluate the level of nucleotide change achieved, residual exonic disruption remaining, globin RNA transcript levels altered, and hemoglobin restored.

With optimized editing reagents and low-density culturing, we demonstrate clinically relevant HDR *in vitro* (E6optE ssODN: 37.9% and rAAV6: 27.4%, E6V ssODN: 38.8% and rAAV6 38%, Figure 1B) and *in vivo* (E6optE BM: 17.7% with ssODN and 10.8% with rAAV6; E6V BM: 15.2% with ssODN and 13% with rAAV6, Figure 2C) using both ssODN and rAAV6 donor template delivery methods following Crispr/Cas9-mediated nuclease disruption. We evaluated both the introduction of a codon-optimized change that leads to the restoration of HbA after RNP-mediated disruption with the E6optE constructs and an introduction of the sickle mutation that leads to sickle globin expression with the E6V constructs.

We observed that human mPBSCs modified with a donor-template had a 1.1 - 1.3-fold decrease in average hCD45^+^ engraftment compared to mock-edited or RNP-edited cells (Figure 2B) ^16, 17, 27^. Given that Mock-edited and RNP-edited cohort did not exhibit a decrease in overall hCD45^+^ engraftment, *ex vivo* manipulation with donor-template delivery seems to have an effect in reducing the overall human chimerism in the BM. This reduction in chimerism occurs irrespective of the mode of donor template delivery and could possibly due to the alteration of the fate/survival or perturbance of quiescence of HSCs due to the presence of an external donor template. It has been reported by other groups that highly specific nuclease reagents can cause transient increase in p53 activation and some delayed proliferation with minimal effect on their functionality. But combining highly-specific nuclease with a donor template possibly causes pronounced proliferative disadvantage, increased p53 activation and cell cycle delays that possibly could alter the fate of these HDR-modified cells ^27, 28^. Further, the donor template modified HSCs could be detected by innate immune sensors in the HSCs, resulting in an inflammatory response that can produce a paracrine effect on the other edited HSCs further causing an amplification of the proliferative disadvantage ^29, 30^. These hypotheses could possibly explain the fact that RNP-edited cells had no deficit in engraftment whereas the combination of nuclease and donor template manipulation results in a deficit in human chimerism in the BM at 16-17 weeks (Figure 2B).

The BM of the animals had comparable levels of multi-lineage engraftment of B-(CD19^+^), myeloid-(CD33^+^) and erythroid-cells (CD235^+^). Although not significant, some level of myeloid skewing was observed with the ssODN-modified animals and some level of lymphoid skewing was observed with the rAAV6-modified animals (Figure S3E-F). The CD34^+^ compartment in the BM was comparable across groups (Figure S3A-C).

The average gene-modified cells within the CD34^+^- and CD235^+^- enriched BM pools were higher in the ssODN-modified animals compared to the rAAV6-modified animals with both E6optE and E6V cohorts (CD34^+^ HDR: ssODN: 25.1% and rAAV6: 9.6%; CD235^+^ HDR: ssODN: 15.2% and rAAV6: 7.6%, Figure 3A). This pattern of higher levels of gene-modified cells with ssODN-edited animals compared to the rAAV6 animals was also observed in the spleen (Figure 3D). A 1.71-fold increase in HDR-modified cells was observed with E6optE ssODN-modified animals and 1.49-fold increase was measured with the E6V ssODN-modified animals compared to the rAAV6 animals in the spleen (Figure 3D).

The globin RNA transcript levels measured in BM at the time of sac further validates the ability of the ssODN donor template to outperform in comparison to the rAAV6 donor template. Though not significant, the average HDR RNA levels measured in individual BMs as well as CD34^+^ and CD235^+^ enriched compartments of BM pools were higher in the ssODN-modified animals than the rAAV6-modified animals (Figure 4A-C, S8A-D).

Functional evaluation of the gene modification in the ex vivo differentiated erythroid cells in the individual BMs as well as the CD34^+^ and CD235^+^ enriched BM pools through IEC identify that Crispr/Cas9-mediated disruption of codon 6 at exon 1 of *HBB* results in a reduction of HbA and an increase in HbF compared to the mock-edited cells (Figure 4A, 4C, S7A-D). The increase in HBF with RNP-mediated disruption could possibly be the result of re-capitulating naturally occurring deletional hereditary persistence of fetal hemoglobin (HPFH) phenotype that produces a beneficial functional outcome resulting in an increase in HbF ^31, 32^.

The HDR event following nuclease disruption with E6optE templates results in the restoration of HbA and with E6V templates results in an increase in sickle globin expression. The ssODN-modified E6optE differentiated BM erythroid cells produced a 1.7-fold increase in HbA compared to RNP-edited cells, whereas the rAAV6-edited cells produced a 1.47-fold increase in HbA compared to RNP-edited cells (Figure 4A). On evaluating the E6V constructs, an average of 21.8% sickle globin was measured in the E6V ssODN-modified animals and an average of 17.2% sickle globin was measured in the rAAV6-modified *ex vivo* differentiated BM erythroid cells (Figure 4B). The HDR-specific globin expression is further accompanied by a 1.6 - 3.4-fold increase in HbF which could prove to be a desirable outcome in the context of a SCD patient cells ^25, 26^. With an increase in residual NHEJ in the BM, higher levels of HBG RNA and HbF protein expression were observed. The persistence of the NHEJ alleles in the BM resulting in an increase in HbF expression is possibly a desirable outcome that seems to persist at 16-17 weeks (Figure S7A-B).

On measuring the β-like globin expression in *ex vivo* differentiated BM erythroid cells as well as the CD34^+^ / CD235^+^ enriched compartment, similar levels of HbA was measured in the E6optE ssODN-modified animals and the rAAV6-modified animals (Figure 4A, S7A, S7C). But, in the *ex vivo* differentiated E6V BM, CD34^+-^, CD235^+^-enriched compartments of BM pools, ssODN-modified animals outperformed rAAV6-modified animals with respect to significantly higher sickle globin expression in the differentiated erythroid cells (Figure 4B, S7B, S7D).

Although several groups have worked on targeted gene editing at the sickle locus ^7, 11-17, 33^, our data compares the method of donor template delivery and its role in altering the *in vivo* persistence of gene modified cells in the BM of NBSGW mice at 16-17 weeks. Clearly both donor template delivery methods are amenable to clinical translation. But our studies demonstrate that the ssODN donor template delivery drives higher sustained engraftment of HDR-modified cells in the BM and HSC compartments which correlates with higher HDR-specific RNA in the BM and HDR-based globin protein expression in the erythroid compartments.

## Materials and Methods

### Culturing and editing CD34^+^ HSC

Chemically-modified sgRNA (Synthego Inc., CA and IDT, Coralville, IA) were designed to cut at codon 6, exon 1 of *HBB*. The optimization of sgRNA was previously reported ^17^. Human mobilized peripheral blood CD34+ cells (hPBSC) were purchased frozen from Cooperative Centers of Excellence in Hematology (CCEH) at Fred Hutchinson Cancer Research Institute (Seattle, WA). CD34^+^ cells were thawed and cultured in SFEM II media with 1µM of SR1, 35 nM UM171 ((STEMCELL technologies, Vancouver, Canada), 100 ng/ml each of hFLT-3L ligand, hTPO, hSCF and hIL-6 (Peprotech, Rocky Hill, NJ). The cells were cultured at low density (0.1 - 0.25×10^6^ cells/ml) ^34^and edited 48 hours post-thaw. RNP mixture was made by combining 20 pmol of Cas9 (*Spy*Fi^™^ Cas9, Aldevron, Fargo, ND) with 50 pmol of sgRNA at room temperature for 15 minutes prior to nucleofection. Five million cells and the RNP mixture were combined in Lonza cuvettes in a total volume of 100 µl P3 buffer and nucleofected using CX100 program (LONZA, Basel, Switzerland). The cells were plated at a density of 1 × 10^6^ cells / ml of SFEM-II after 5 minutes in recovery media. Chemically modified ssODN (either E6optE GAA or E6V GTC) 50 pmol / 2 × 10^5^ cells were delivered along with the RNP mixture at the time of nucleofection ^17^. rAAV6 were designed as previously described ^17^. The rAAV6 was diluted to 1% culture volume (GTC rAAV6 ∼MOI of 901, GAA rAAV6 ∼MOI of 2,190) in SFEM II media and added to RNP-nucleofected cells. The rAA6 media was removed 16-18 hours post-transduction. 2 × 10^5^ cells from each group were maintained in erythroid differentiation media (IMDM media (Fisher Scientific, Hampton, NH) containing 1 ng/ml hIL-3, 2 IU/ml EPO, 20 ng/ml h-SCF, 20% heat-inactivated FBS and 1% Pen/Strep (PeproTech, Rocky Hill, NJ) for 14 days to measure HDR, RNA expression and Globin expression.

### Repair templates

The ssODN and rAAV6 were designed as previously reported ^17^. ssODNs were purchased from Integrated DNA Technologies. The ssODNs E6V GTC introducing the sickle mutation and E6optE GAA introducing the codon-optimized change were 168 bp long with 2 terminal nucleotides on the 5’ and 3’ ends modified by phosphorothioate bonds. The rAAV6 E6V GTC introducing the sickle mutation and E6optE rAAV6 introducing the codon-optimized change at codon 6 of exon 1 of *HBB* were cloned with 2.2 Kb homology arm and prepared according to previously described protocol ^35^. The rAAV6 vector, serotype helper and HgT1-adeno helper plasmids ^36^ were transfected into HEK293T cells and harvested at 48 hours and treated with benzonase. An iodixanol density gradient was used to purify the virions with recombinant rAAV6 genomes. The qPCR-based titers of AAV genomes were determined by using ITR specific primers and probe ^37^.

### Engraftment studies in NBSGW mice

NBSGW mice were purchased from Jackson Laboratories. These mice were maintained according to the Association for Assessment and Accreditation of Laboratory Animal Care (AAALAC) standards by the Seattle Children’s Research Institute (SCRI) with the approval of SCRI Institutional Animal Care and Use Committee (IACUC). Mice between 6-8 weeks of age were treated with 12.5 mg/kg of busulfan (Selleckchem) 24 hours pre transplant, and Baytril® (Bayer, Shawnee Mission, KS) was given 48 hours pre transplant. At time of transplant 1.6 - 2 × 10^6^ edited cells were delivered by tail vein injection 24 hours after Busulfan. Animals in the mock-edited group were given the same number of cells that had been cultured in the identical conditions but were not nucleofected. BM and spleen were harvested from mice at 16-18 weeks post-transplant and were analyzed by FACS for hCD45^+^, mCD45^+^ as well as multi-lineage engraftment of CD19^+^, CD33^+^, CD235^+^, CD3^+^, CD34^+^, CD38^+^ cells (Table S3). BM and spleen were harvested for gDNA and RNA extraction. One million cells from each individual mouse BM were cultured in erythropoietin containing differentiation media and harvested after 14 days of *ex vivo* culturing. Cells were kept below a million/ml density to avoid overcrowding.

### BM enrichment of CD34+ and CD235 compartments

BM within each group were pooled and enriched with CD34+ beads (Miltenyi Biotec, Gladbach, Germany). The bound fraction was collected and eluted in MACS buffer according to manufacturer’s protocol. The flow through from the column called the CD34^-^ was used for enriching the CD235^+^ compartment using GP235+ beads (Miltenyi Biotec, Gladbach, Germany). The enriched CD34+ and CD235+ cells were collected, checked for purity and then pelleted for gDNA, RNA and ex vivo differentiation.

### Measuring HDR events using ddPCR

DNA from bone marrow and spleen were analyzed according to previous protocol ^17^. DNeasy blood and tissue kit (Qiagen, Venlo, Netherlands) was used to extract gDNA, samples were RNase-treated. A ddPCR F/R primers were used to amplify a 210 bp amplicon with in HBB from 50-100 ng of gDNA after digestion with ECORV-HF that shears the gDNA outside of the amplicon region. The assay was designed as a dual probe assay with a wild type HEX (WT) and HDR-FAM probe competing for overlapping binding sites in the same reaction (Table S1-S2). The reference HEX probe was run in parallel in a separate well with the same primers. The ddPCR reaction was prepared with ddPCR supermix for probes (No dUTP) and droplets were generated on the QX200 droplet generator (Bio-rad, Hercules, CA). The PCR was run using a Bio-RAD PCR machine and the fluorescence intensity was measured using QX200 droplet reader. The % FAM^+^ (HDR) and HEX^+^ (WT) events were calculated after correction with reference HEX^+^ events. .

### RNA transcript analysis by ddPCR

Cells were collected for RNA and resuspended in 350 ul of RLT buffer before freezing at -80°C. RNA was extracted using RNeasy purification kit (Qiagen, Venlo, Netherlands). after DNase treatment using the manufacturer’s protocol. 30-40 ng of RNA was used to make cDNA using the iScript Adv cDNA Syn Kit (Bio-Rad life Sciences). The RNA was diluted 1:50 in water for ddPCR analysis. A 210 bp region was amplified using ddPCR F/R primersand a FAM^+^ HDR and HEX^+^ WT probe (Table S1-S2). In a parallel reaction GAPDH primers along with GAPDH-HEX^+^ probes were used as a reference to amplify a 196 bp amplicon. The same reaction had a total HBG FAM probe with HBG primers to detect change in total HBG as a result of editing. HDR RNA, *HBB* RNA and *HBG* RNA were measured after adjusting for *GAPDH* RNA as a reference. The RNA transcripts were quantified using the following equation:

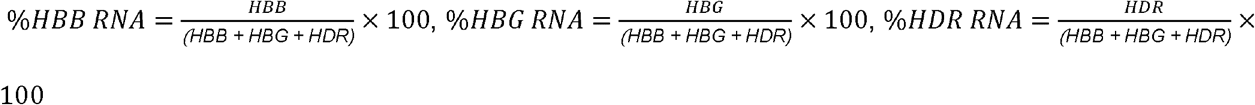

### Measuring INDEL frequencies

gDNA was amplified around the cut site with forward and reverse HBB-1250 F/R primers. The PCR products were column purified using Machery Nagel kit (Düren, Germany) and 50-100 ng of purified PCR product were sent to Genewiz (South Plainfield, NJ) with the sequencing primer SCL-386 reverse primer. The sequences were analyzed using the ICE algorithm (Synthego Inc., CA) to measure the amount of HDR and NHEJ following editing. The algorithm reports ICE score and Knock-in (KI-Score). We calculated NHEJ = ICE score-KI score.

### HPLC analysis of edited cells

Cells from the erythroid differentiation media were washed in PBS and frozen at -80°C. The cells were lysed right before analysis in HPLC-grade water and sent for analysis at Fred Hutchinson Cancer Research Institute (Seattle, WA). *In vitro* cultures of edited cells were analyzed using reverse phase (RP-HPLC) and/or ion exchange (IEC) chromatography.

### IEC Analysis

All the cells from transplant experiments were analyzed using IEC to measure HbA, HbA2, HbS, and HbF. For Ion Exchange Chromatography (IEC), samples were provided in HPLC-grade water and kept frozen at -80°C until just prior to analysis. The analysis was conducted on a Thermo Scientific Vanquish Horizon UHPLC connected to a diode array UV detector with a 10mm pathlength light-pipe flow cell. Samples were gradient-eluted from a PolyCAT-A 200×2.1mm 5µm, 1000Å column (PolyLC p/n 202CT0510) using a gradient of 2-100% “B” over a 24-minute period at a flowrate of 300µL/minute. The HPLC mobile phases consisted of “A” = 40mM Tris/3mM KCN in HPLC-grade water adjusted to pH 6.4 with concentrated acetic acid and “B” of the same composition with the addition of 200mM NaCl. The column was held at 30°C throughout the run, and the autosampler compartment was maintained at 4°C. The detected signal at 418nm was recorded in Chromeleon (ver 7). A “blank” run followed by a reference standard run were acquired prior to running samples. The reference standard was a 5µL injection of a mixture of HbA, HbF, and HbS (at 0.5µg/µL, 1.0µg/µL, and 0.5µg/µL respectively) and was used to establish the elution times of each globin tetramer.

### RP-HPLC Analysis

For Reverse-phase (RP) chromatography, samples were removed from -80°C, allowed to thaw on ice, and lysed by adding approximately 70uL of HPLC-grade water. The samples were then centrifuged at 20,000x g for 10 minutes at 4°C and the supernatant was transferred to a chromatography vial for analysis. Following lysis, samples were gradient-eluted from an Aeris Widepore C4 2.1×250mm 3.6µm 200Å column (Phenomenex p/n 00G-4486-AN) at a flowrate of 150µL/minute. The gradient used was as follows: time 0: 37% “B” followed by holding at 37% “B” for 1 minute, then ramping up to 55% “B” in 60 minutes, holding at 55% “B” for 5 minutes, then ramping down to 37% “B” over 2 minutes, and then finally re-equilibrating the column at starting conditions for 23 minutes. The mobile phases consisted of “A” = H_2_O/trifluoroacetic acid (TFA) at 0.1% (by volume), and “B” = acetonitrile/methanol (80:20 v:v) and TFA at 0.08%. The column temperature was held at 45°C and the autosampler compartment was maintained at 4°C. The detected peaks at 205nm were recorded in Chromeleon. Similar to the IEC method, a blank and reference standard run were acquired prior to running samples. The reference standard made from a mix of adult, fetal and sickle hemoglobin (0.5µg/uL each) was used to establish elution times of the globin monomers using a 10µL injection.

### Statistical Analysis

The data collected from experiments were analyzed on Graph Pad Prism 7 using one-way or two-way ANOVA analysis with Dunnett’s multiple comparisons test. All samples were compared to Mock-treated cells to evaluate the significance, * *p* < 0.05 ** *p* < 0.01 *** *p* < 0.001 ****, *p* <0.0001.

## Author Contributions

Conceptualization S.P., D.J.R., Methodology, S.P., D.J.R.,

Investigation S.P., S.N.L, M.P.B., Writing S.P., D.J.R., Original Draft S.P., Writing

Review & Editing S.P., D.J.R., Funding Acquisition D.J.R., Resources, D.J.R, Supervision S.P., D.J.R,

## Acknowledgements

We would like to thank Claire Stoffers, Ezra Lopez and Stefan Lachkar for their help with animal experiments. We would like to thank the Seattle Children’s Research viral core members; Iram F. Khan and Christopher Zavala Galvan for making all the rAAV6 viruses and Brian P. Milless from the proteomics core, Fred Hutchinson Cancer Research Institute for optimizing all the HPLC/IEC runs. We would like to thank Dr. Christopher T. Lux for scientific discussions and his help with setting up the methocult assay.

## Conflict of interest

The authors declare that they have nothing to disclose.

**Supplemental Table 1.**
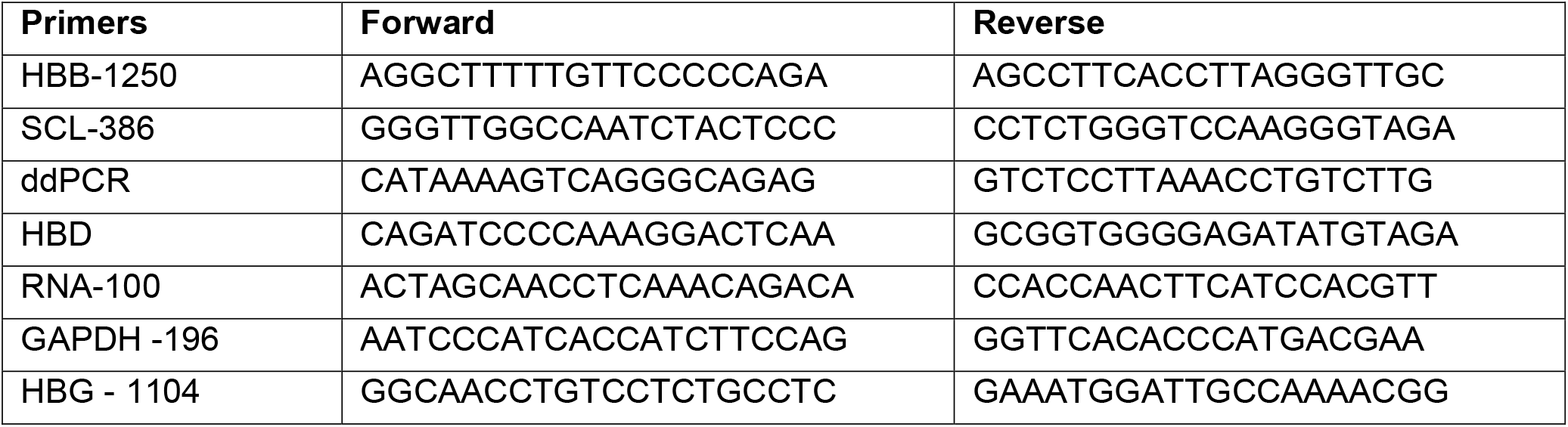
Primers used for analysis.

**Supplemental Table 2.**
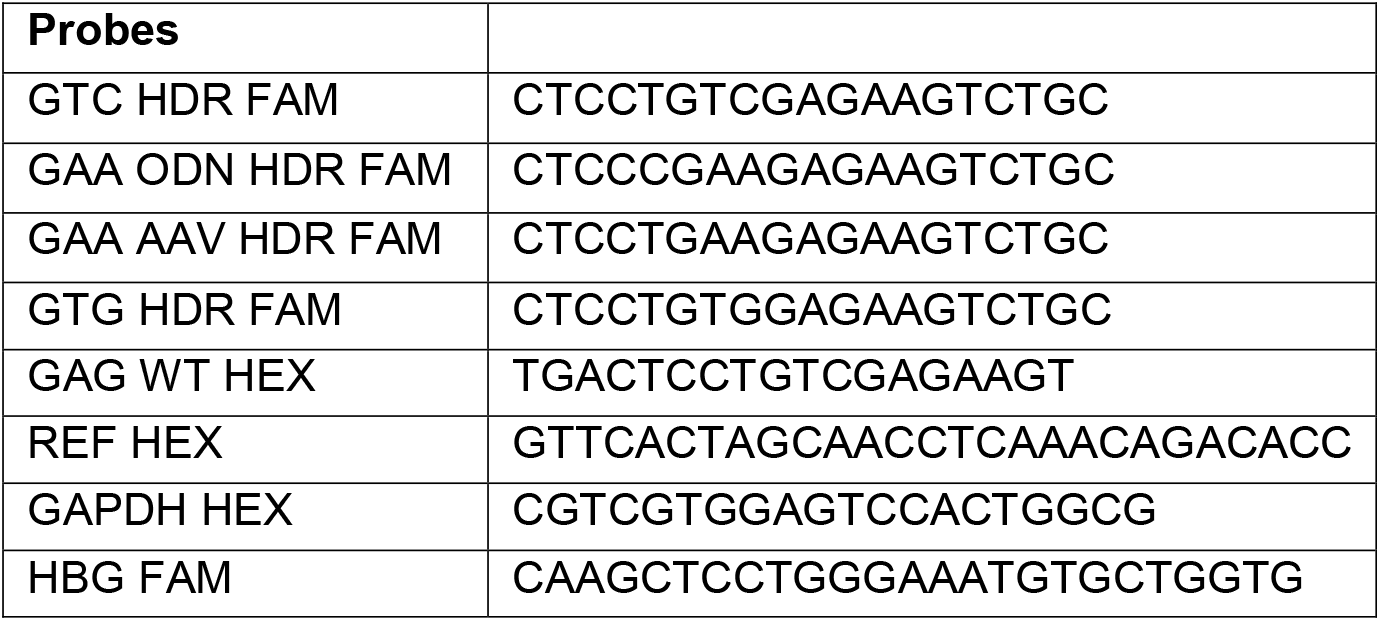
Probes used for analysis.

**Supplemental Table 3.**
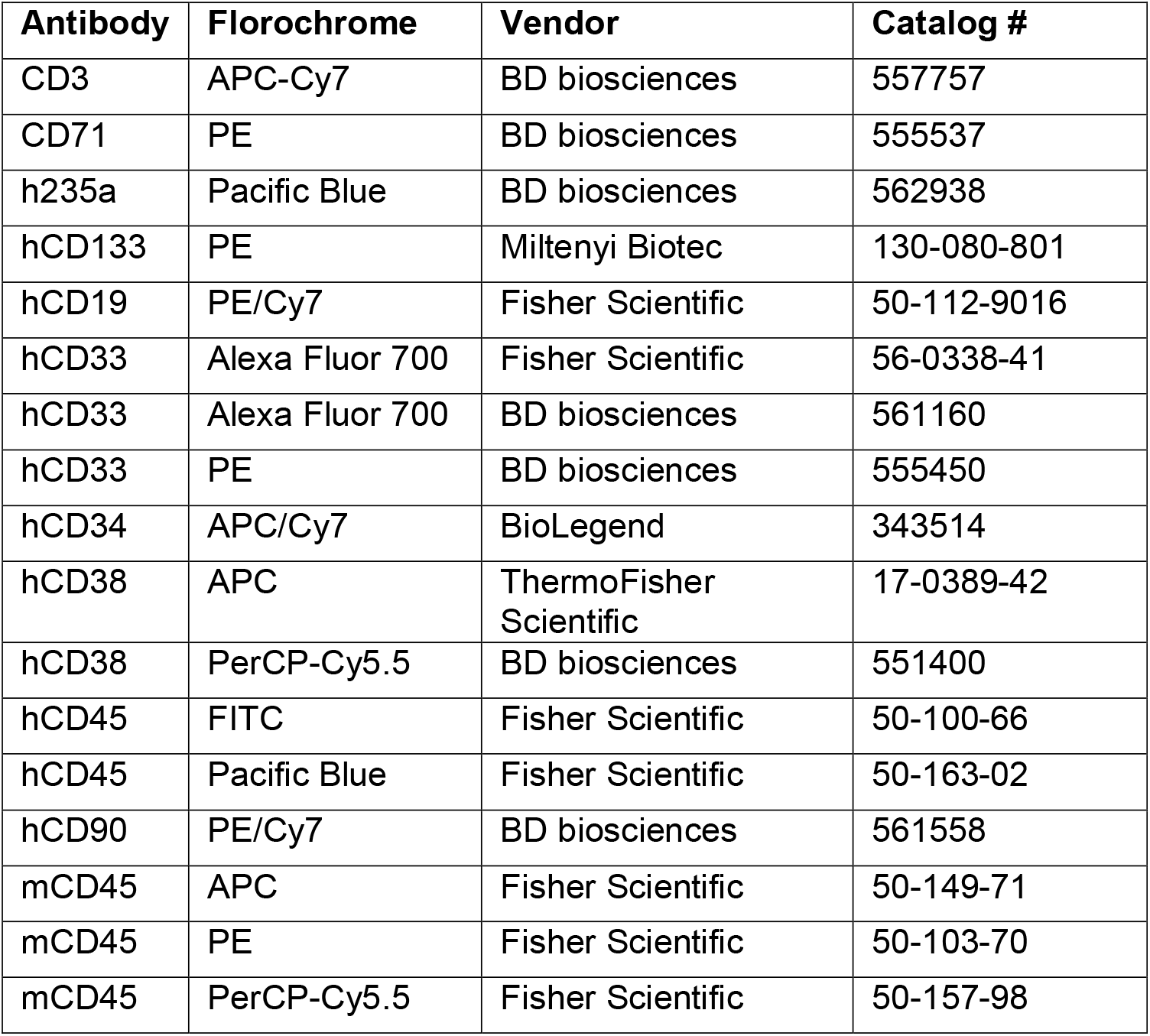
List of antibodies used for analysis.

| **Antibody** | **Fluorochrome** | **Vendor** | **Catalog #** |
| --- | --- | --- | --- |
| CD3 | APC-Cy7 | BD biosciences | 557757 |
| CD71 | PE | BD biosciences | 555537 |
| h235a | Pacific Blue | BD biosciences | 562938 |
| hCD133 | PE | Miltenyi Biotec | 130-080-801 |
| hCD19 | PE/Cy7 | Fisher Scientific | 50-112-9016 |
| hCD33 | Alexa Fluor 700 | Fisher Scientific | 56-0338-41 |
| hCD33 | Alexa Fluor 700 | BD biosciences | 561160 |
| hCD33 | PE | BD biosciences | 555450 |
| hCD34 | APC/Cy7 | BioLegend | 343514 |
| hCD38 | APC | ThermoFisher Scientific | 17-0389-42 |
| hCD38 | PerCP-Cy5.5 | BD biosciences | 551400 |
| hCD45 | FITC | Fisher Scientific | 50-100-66 |
| hCD45 | Pacific Blue | Fisher Scientific | 50-163-02 |
| hCD90 | PE/Cy7 | BD biosciences | 561558 |
| mCD45 | APC | Fisher Scientific | 50-149-71 |
| mCD45 | PE | Fisher Scientific | 50-103-70 |
| mCD45 | PerCP-Cy5.5 | Fisher Scientific | 50-157-98 |

## Figure Legends

**Figure S1.**
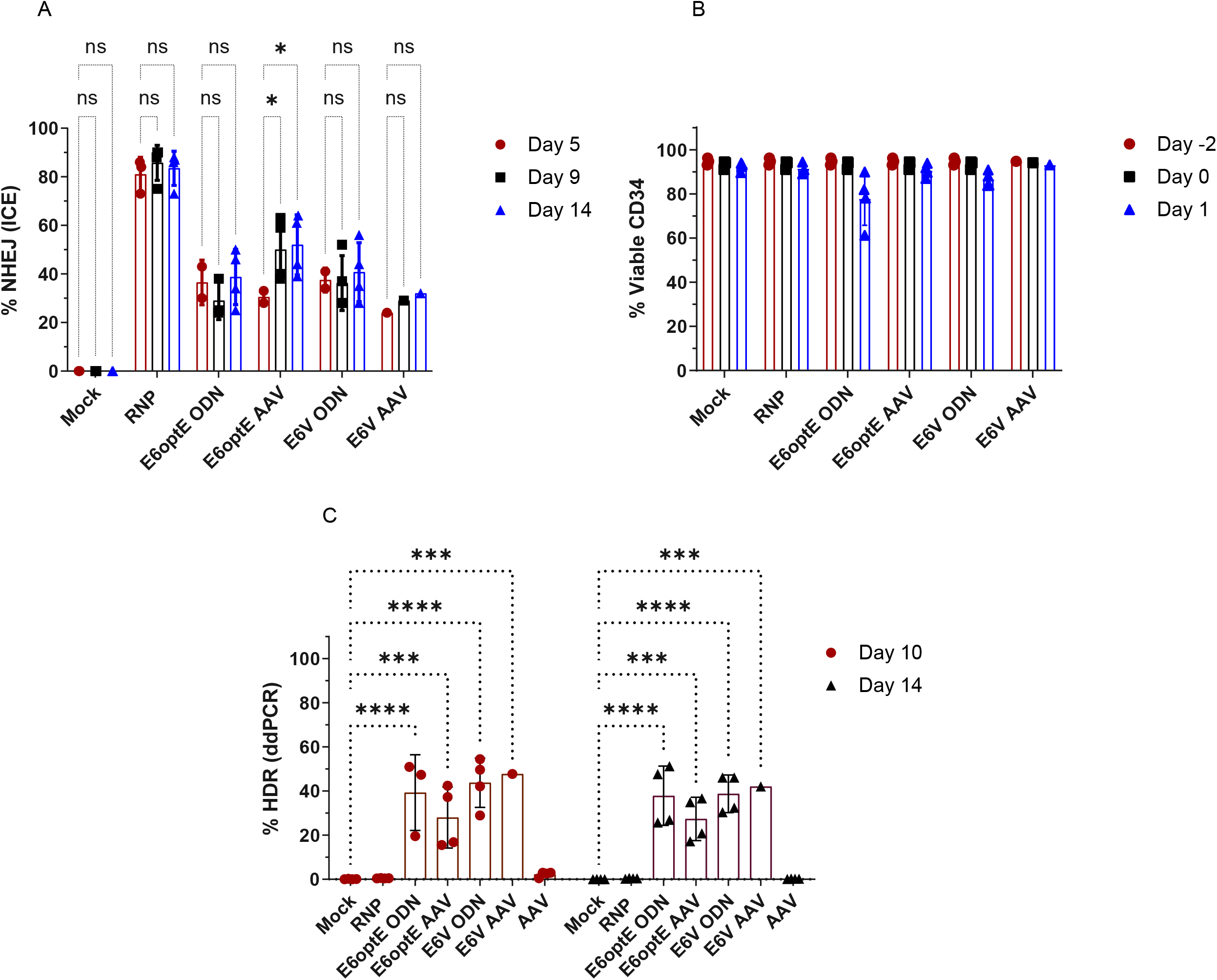
*In vitro* analysis of transplant input cells. (A) The NHEJ % measured by ICE algorithm on day 5, 9, 14 post-editing in *in vitro* cultured input cells. (B) % viability of Mock-edited, RNP-edited, ssODN-modified and rAAV6-modified mPBSC input CD34^+^ cells, on day-2 (Thaw), Day 0 (Editing) Day 2 post-editing (n = 4 transplant). (C) The HDR % measured by ddPCR using a dual probe assay (FAM/HEX) on day 10 and 14 post-editing in *in vitro* cultured transplant input cells. Statistical analyses were run using 2-way ANOVA using Tukey’s multiple comparison test (P-value of <0.0001, *** < 0.001, ** <0.01).

**Figure S2.**
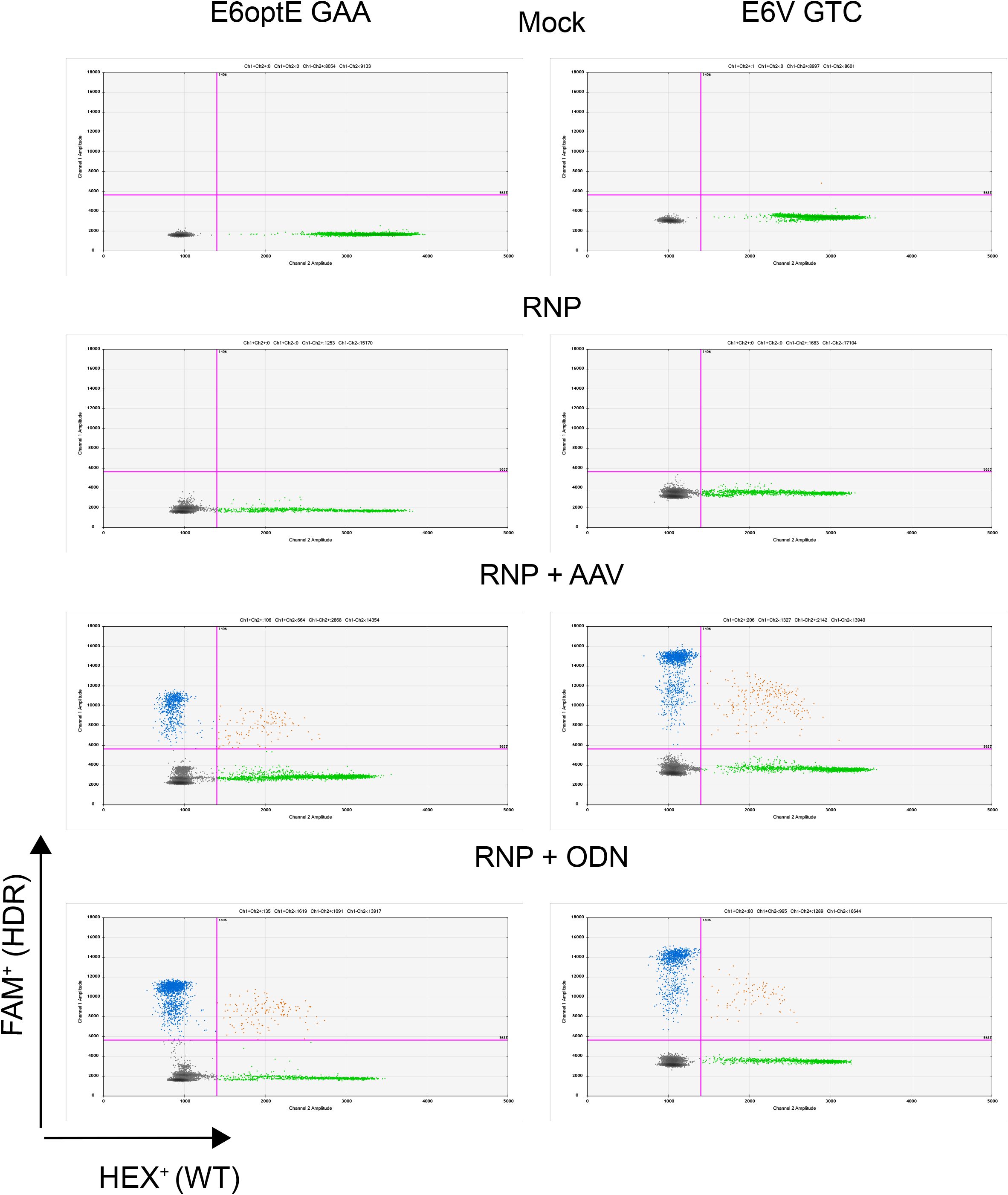
ddPCR assay to measure HDR and WT alleles. HDR% and WT% measured by ddPCR using a dual probe assay (HDR-FAM/WT-HEX). Data shows 2D plots of amplitudes measuring HDR-FAM^+^ and WT-HEX^+^ signals in Mock, RNP, RNP + AAV and RNP +ODN samples within E6optE and E6V cohorts in the extracted gDNA from BM of individual animals at 16-17 weeks.

**Figure S3.**
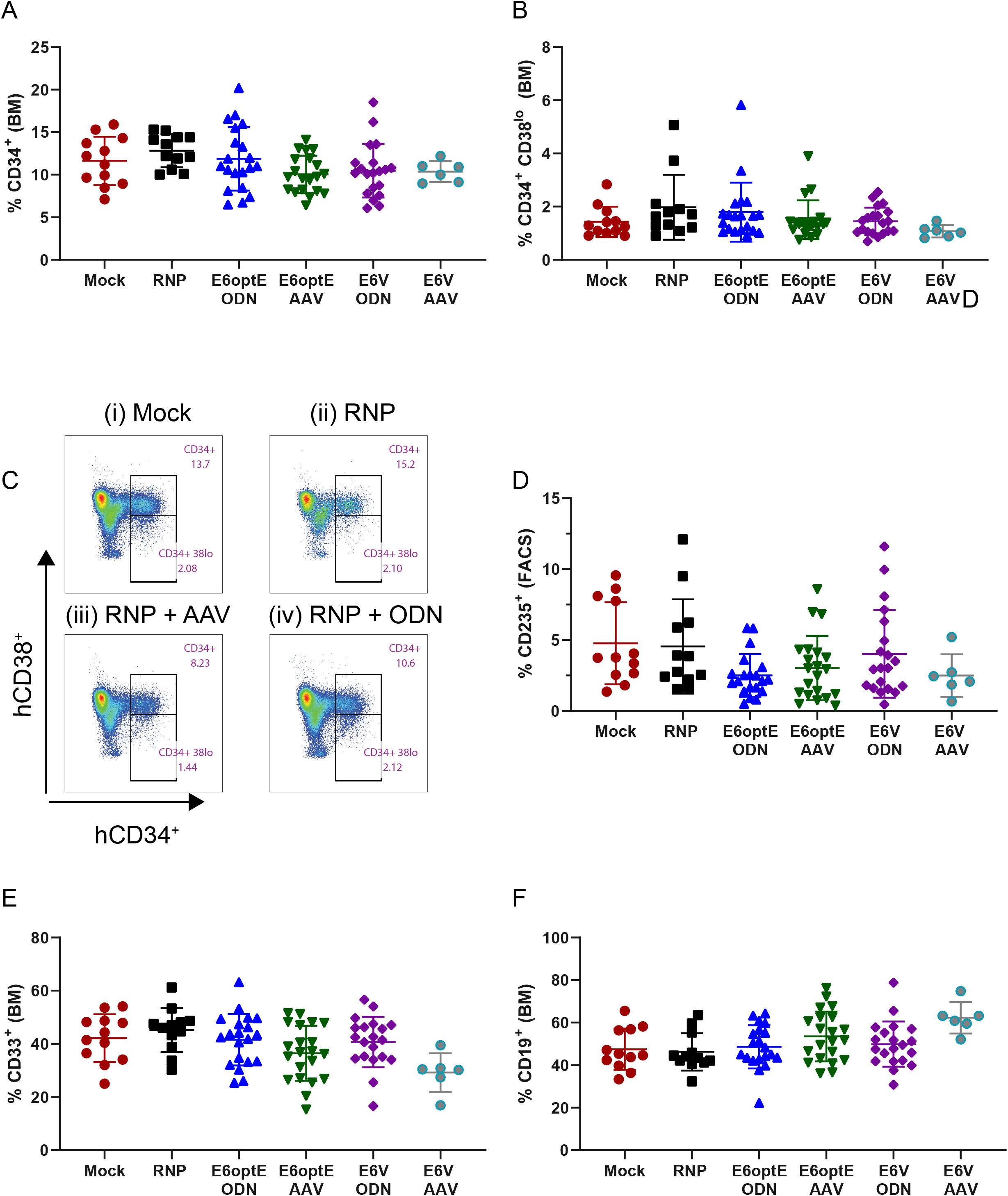
Multi-lineage engraftment in the BM. (A) % hCD34^+^, (B-C) % hCD34^+^ CD38^lo^ populations (D) %CD235^+^ (E) %hCD33^+^ (F) %hCD19^+^ measured in the BM by FACS at 16-17 weeks across mock-edited, RNP-edited, E6optE GAA ssODN/rAAV6-modified, E6V GTC ssODN/rAAV6-modified cohorts. Gating strategy: Live > single cells > hCD45^+^ > hCD34^+^ CD38^lo^populations. Statistical analyses were run using 2-way ANOVA using Tukey’s multiple comparison test (P-value of <0.0001, *** < 0.001, ** <0.01). All comparisons were non-significant.

**Figure S4.**
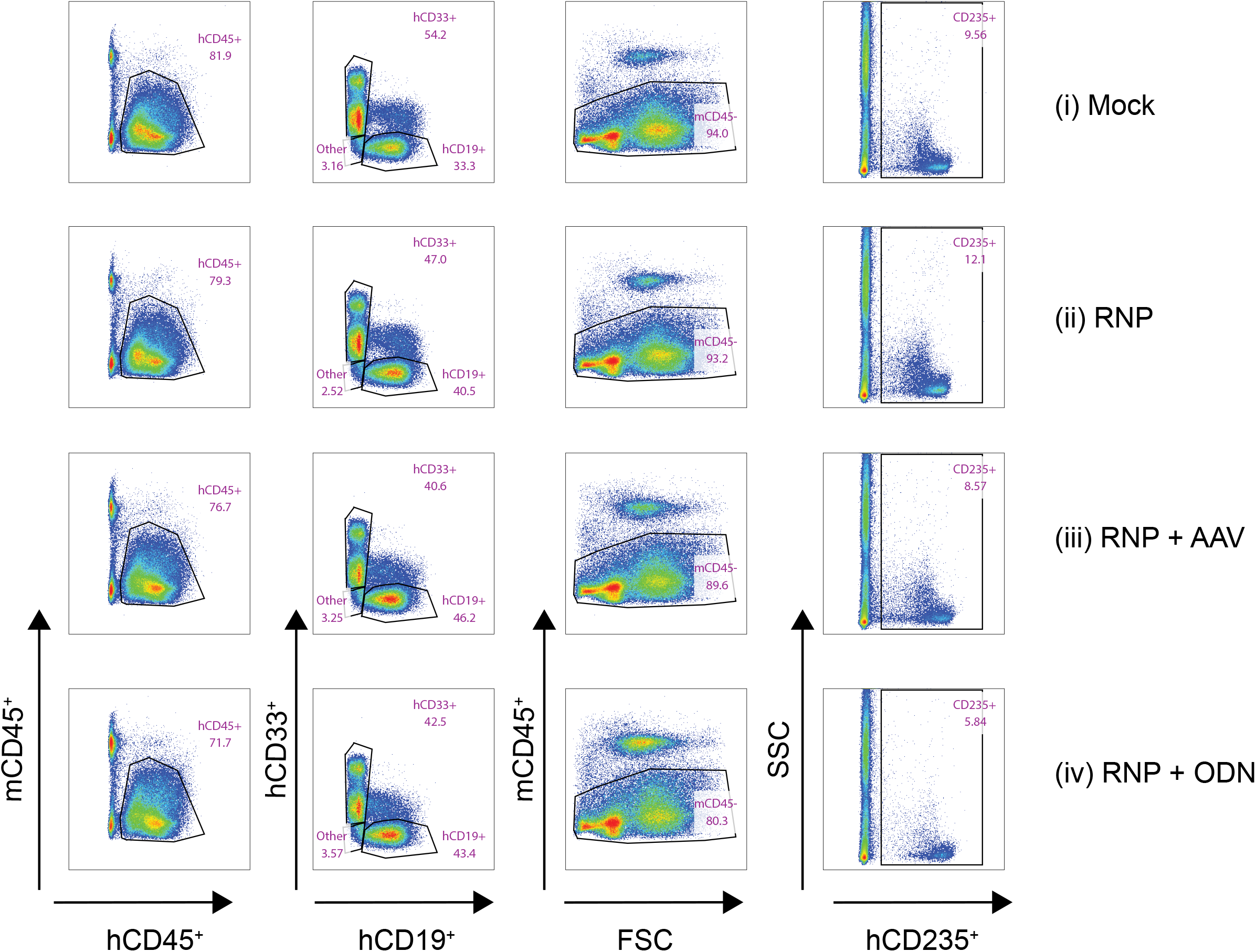
Flow cytometry analysis and multi-lineage engraftment. Representative flow plot of Mock-edited, RNP-edited, E6optE GAA ssODN/rAAV6: hCD45^+^, hCD19^+^, hCD33^+^ and hCD235^+^ populations in the BM of NBSGW mice across Mock-edited, RNP-edited . Gating strategy: Live > single cells > hCD45^+^ > hCD19^+^/hCD33^+^ and Live > single cells > mCD45^-^ > hCD235^+^.

**Figure S5.**
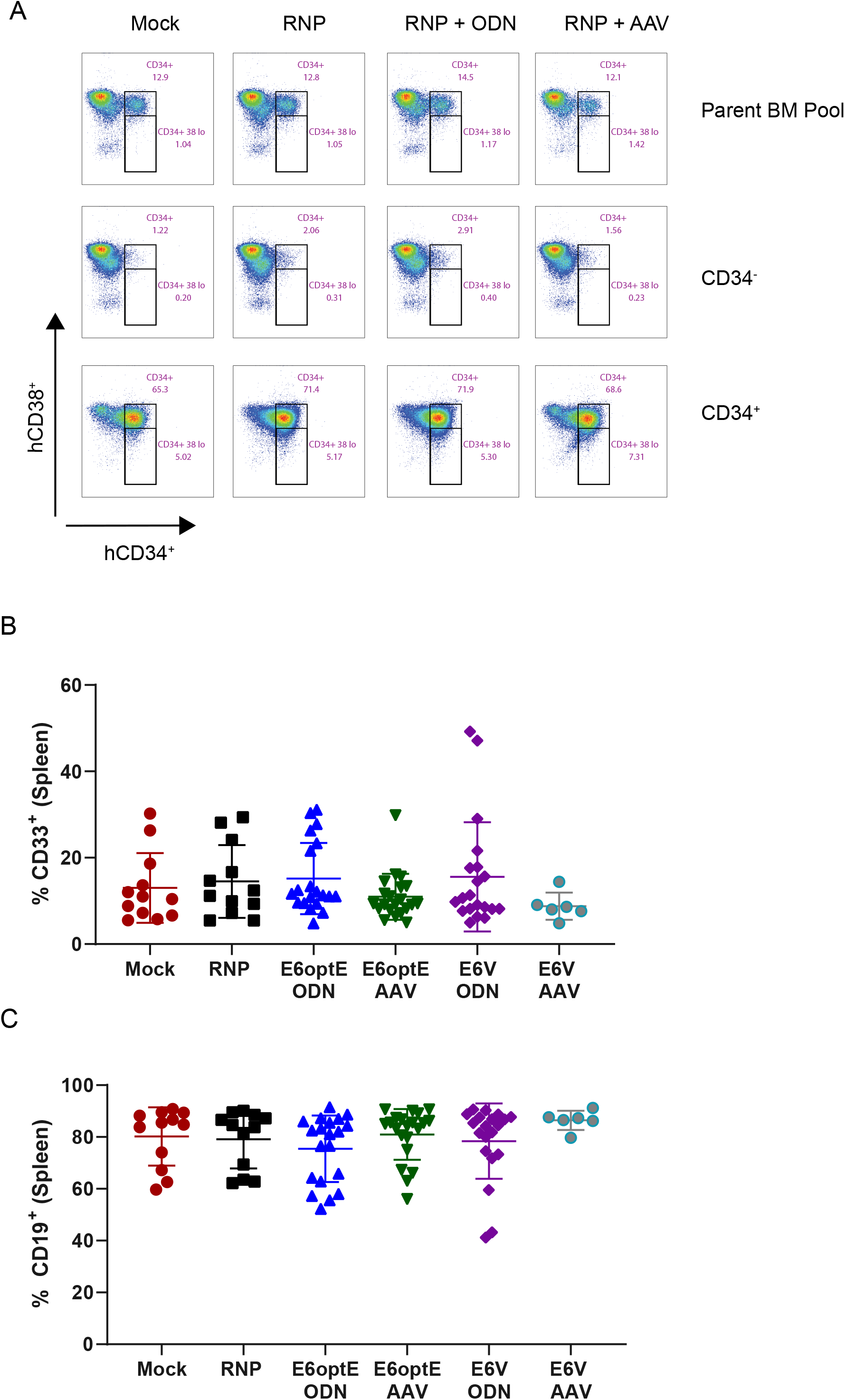
hCD34^+^ enrichment and multi-lineage engraftment in the Spleen. (A) Representative flow plot of hCD34^+^ enrichment using magnetic beads from BM pools in Mock-edited, RNP-edited, E6optE GAA ssODN/rAAV6 groups. Top panel shows parent BM pools, middle panel CD34^-^ fractions, bottom panel: CD34^+^ fractions. Gating strategy used: Live > single cells > hCD45^+^ > hCD34^+^. (B) %hCD33^+^ (C) %hCD19^+^ measured in the spleen across mock-edited, RNP-edited, E6optE ssODN/rAAV6-modified, E6V ssODN/rAAV6-modified cohorts. Gating strategy: Live > single cells > hCD45^+^ > hCD19^+^/hCD33^+^. Statistical analyses were run using 2-way ANOVA using Tukey’s multiple comparison test (P-value of <0.0001, *** < 0.001, ** <0.01).

**Figure S6.**
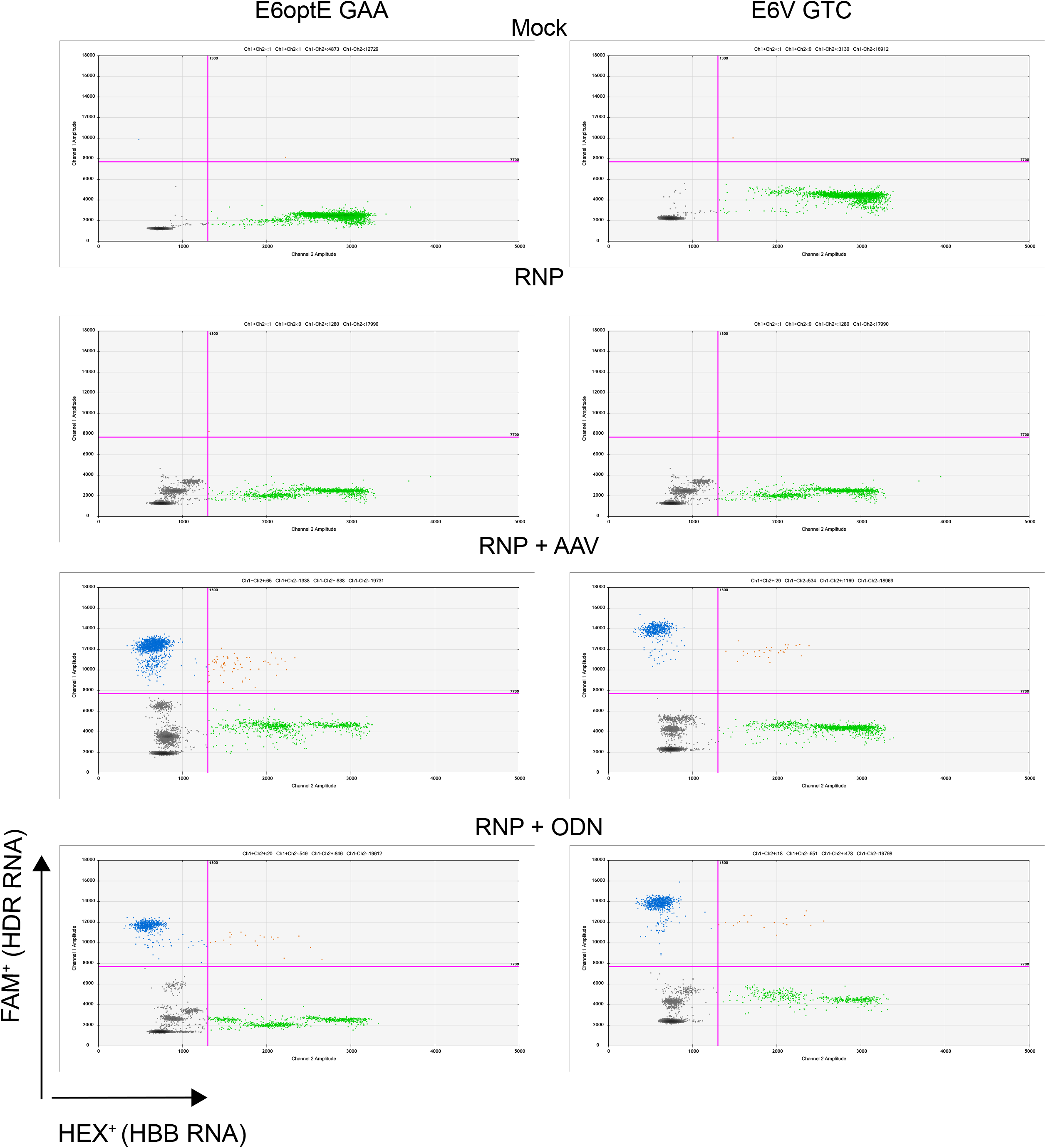
ddPCR assay to analyze the BM RNA transcript levels. Representative ddPCR populations measuring FAM^+^ HDR RNA levels and HEX^+^ *HBB* WT RNA levels in purified total RNA from the BM of individual animals at 16-17 weeks across mock-edited, RNP-edited, E6optE ssODN/rAAV6-modified, E6V ssODN/rAAV6-modified cohorts.

**Figure S7.**
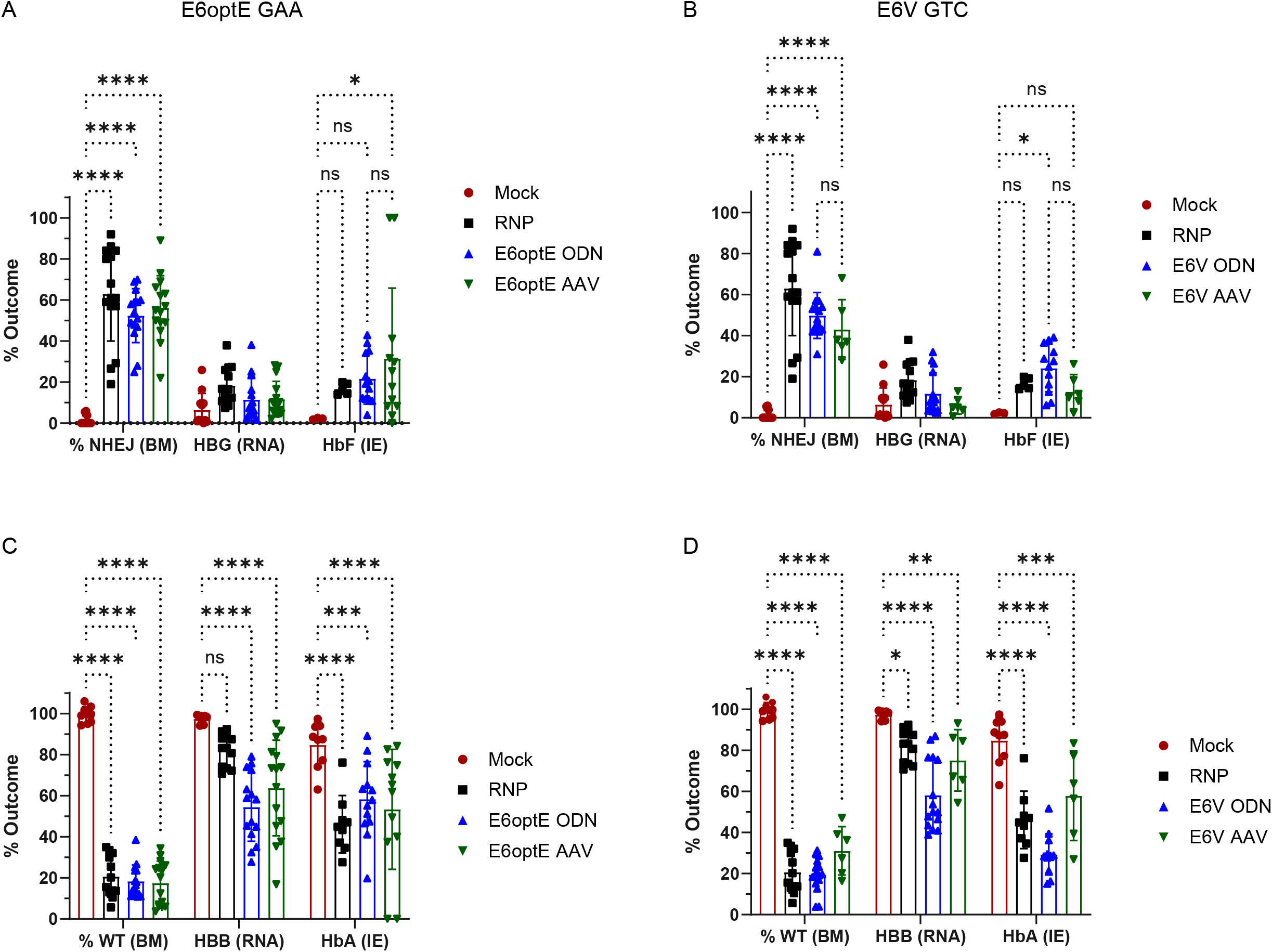
Functional modulation of RNA and hemoglobin in response to NHEJ and WT allele in the BM. NHEJ% and HBG RNA measured in individual BM of animals and HbF protein levels measured in individual *ex vivo* erythroid differentiated BM cultures of (A) E6optE ssODN/rAAV6-modified (B) E6V ssODN/rAAV6-modified cohorts. BM WT% and HBB RNA measured in individual BM of animals and HbA protein levels measured in individual *ex vivo* erythroid differentiated BM cultures of (C) E6optE ssODN/rAAV6-modified (D) E6V ssODN/rAAV6-modified cohorts. Statistical analyses were run using 2-way ANOVA using Tukey’s multiple comparison test (P-value of <0.0001, *** < 0.001, ** <0.01).

**Figure S8.**
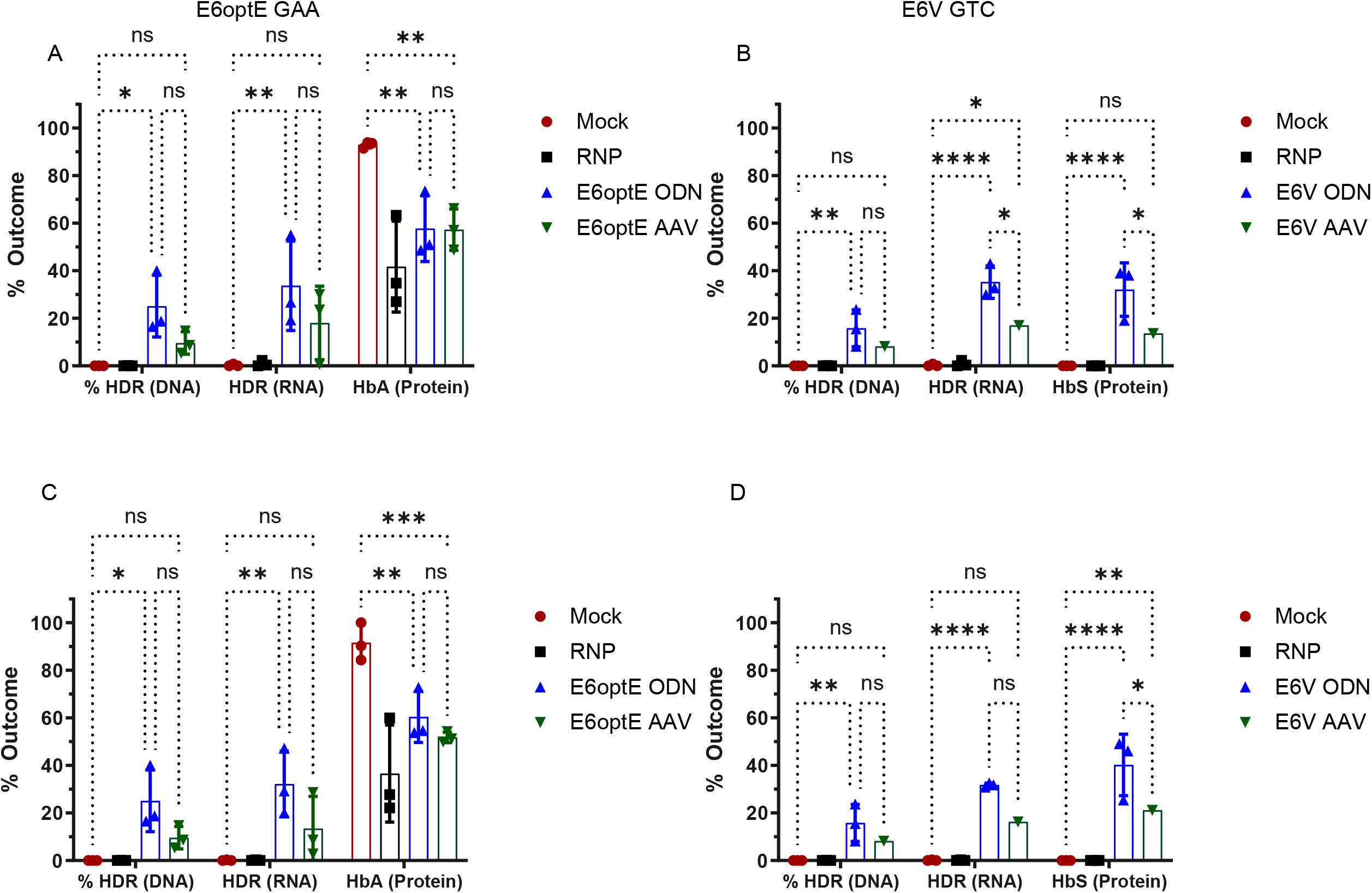
Functional modulation of RNA and hemoglobin in response to HDR within the CD34^+^ and CD235^+^ enriched compartments. HDR% and HDR RNA measured in BM pools and modulation of HbA within the E6optE cohort and HbS within the E6V cohort in *ex vivo* erythroid differentiated CD34^+^ enriched BM pools of (A) Mock-edited, RNP-edited, E6optE ssODN/rAAV6-modified cohorts. (B) Mock-edited, RNP-edited, E6V ssODN/rAAV6-modified cohorts (n = 3 transplants). HDR% and RNA measured in BM pools and modulation of HbA within the E6optE cohort and HbS within the E6V cohort in *ex vivo* erythroid differentiated CD235^+^ BM pools of (C) Mock-edited, RNP-edited, E6optE ssODN/rAAV6-modified cohorts. (D) Mock-edited, RNP-edited, E6V ssODN/rAAV6-modified cohorts. Statistical analyses were run using 2-way ANOVA using Tukey’s multiple comparison test (P-value of <0.0001, *** < 0.001, ** <0.01).

**Figure S9.**
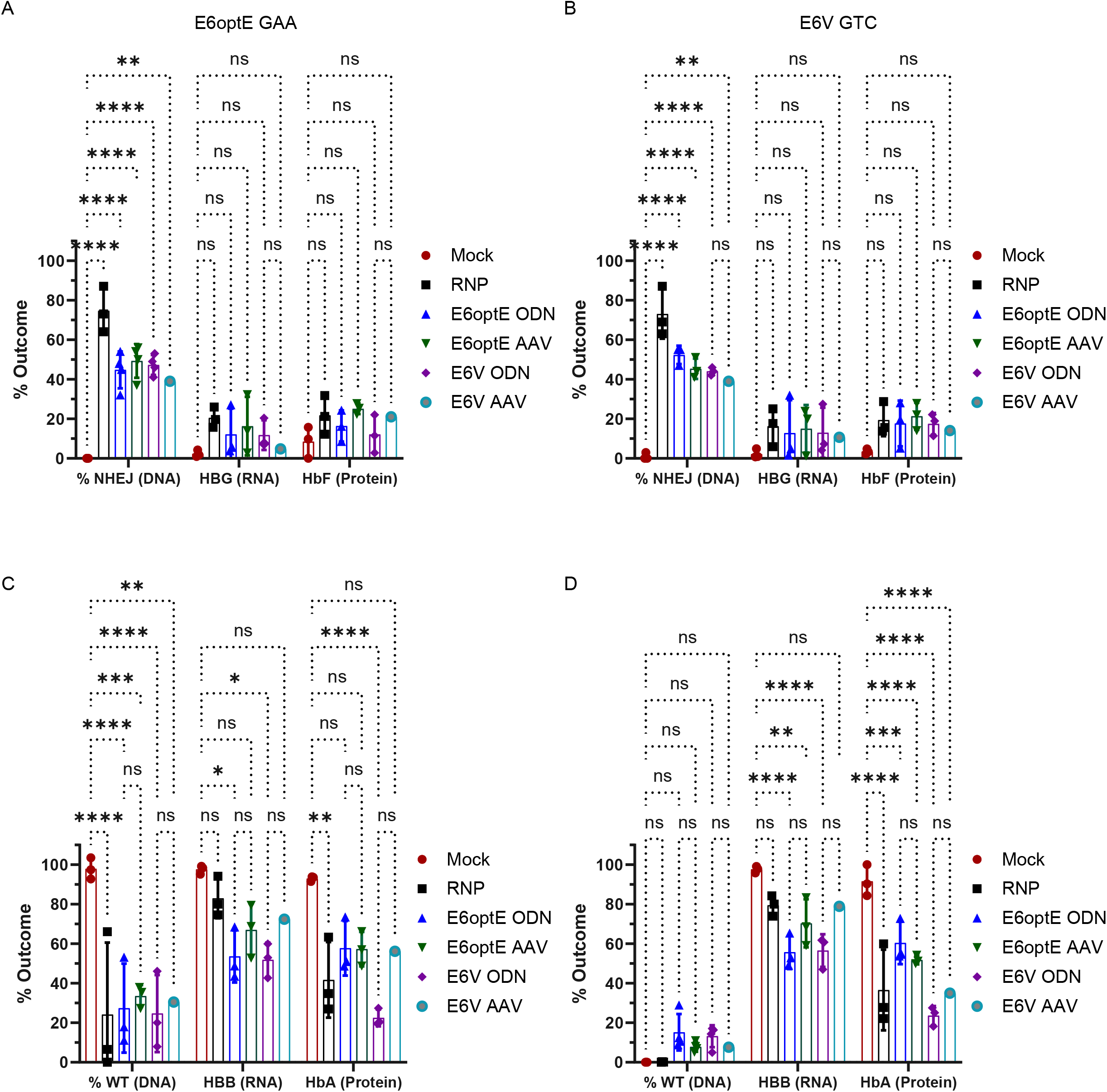
Functional modulation of RNA and hemoglobin in response to NHEJ/WT alleles within the CD34^+^ and CD235^+^ enriched compartments. NHEJ% and HBG RNA measured in BM pools and modulation of HbF in *ex vivo* erythroid differentiated BM pools of Mock-edited, RNP-edited, E6optE and E6V ssODN/rAAV6-modified cohorts measured in (A) CD34^+^ enriched BM pool (B) CD235^+^ enriched BM pools. WT% and HBB RNA measured in BM pools and modulation of HbA in *ex vivo* erythroid differentiated BM pools of Mock-edited, RNP-edited, E6optE and E6V ssODN/rAAV6-modified cohorts measured in (C) CD34^+^ enriched BM pool (D) CD235^+^ enriched BM pools. Statistical analyses were run using 2-way ANOVA using Tukey’s multiple comparison test (P-value of <0.0001, *** < 0.001, ** <0.01).

